# Chemical genetics of AGC-kinases reveals shared targets of Ypk1, Protein Kinase A and Sch9

**DOI:** 10.1101/756841

**Authors:** Michael Plank, Mariya Perepelkina, Markus Müller, Stefania Vaga, Xiaoming Zou, Marina Berti, Jacques Saarbach, Steven Haesendonckx, Nicolas Winssinger, Ruedi Aebersold, Robbie Loewith

## Abstract

Protein phosphorylation cascades play a central role in the regulation of cell growth and protein kinases PKA, Sch9 and Ypk1 take centre stage in regulating this process in *S. cerevisiae*. To understand how these kinases co-ordinately regulate cellular functions we compared the phospho-proteome of exponentially growing cells without and with acute chemical inhibition of PKA, Sch9 and Ypk1. Sites hypo-phosphorylated upon PKA and Sch9 inhibition were preferentially located in RRxS/T-motifs suggesting that many are directly phosphorylated by these enzymes. Interestingly, when inhibiting Ypk1 we not only detected several hypo-phosphorylated sites in the previously reported RxRxxS/T-, but also in an RRxS/T-motif. Validation experiments revealed that neutral trehalase Nth1, a known PKA target, is additionally phosphorylated and activated downstream of Ypk1. Signalling through Ypk1 is therefore more closely related to PKA- and Sch9-signalling than previously appreciated and may perform functions previously only attributed to the latter kinases.

## INTRODUCTION

Cell growth is dynamic and highly regulated by signalling pathways that are conserved across evolution. To accomplish this regulation, eukaryotes have developed intricate means to assess growth conditions and to rapidly communicate this information to the processes controlling the accumulation of mass, the modification of cellular volume and of membrane surface area. Although many signal transduction pathways are involved in this regulation, those employing AGC-family kinases (named after Protein Kinases A, G and C) are prominent.

The target of rapamycin complexes 1 and 2 (TORC1 and TORC2) are central sensors of environmental conditions and regulators of cell growth (1). Both complexes exert their functions by phosphorylating AGC-kinases as their main targets. In *S. cerevisiae* TORC1 primarily responds to changes in carbon and nitrogen availability and regulates ribosome biogenesis, cell cycle progression and stress responses via the AGC-kinase Sch9, similar to S6K downstream of mammalian TORC1 (1, 2).

Another AGC-kinase, protein kinase A (PKA), performs many, if not most of its functions in parallel to Sch9 by regulating an overlapping set of functions and potentially by cross-talk (3–5). Indeed, Sch9 was originally identified by virtue of its ability to suppress growth phenotypes associated with loss of PKA activity (6). Reciprocally, hyper-activation of PKA signalling can suppress phenotypes linked to the loss of Sch9 activity (4).

Tpk1, Tpk2 and Tpk3 are the partially redundant paralogs of the catalytic subunit of PKA. When cells are starved of carbon, cyclic AMP (cAMP) levels are low and, as a consequence, PKA is kept inactive by its regulatory subunit Bcy1. Glucose addition induces activation of the adenylate cyclase Cyr1/Cdc35 via the small GTPase Ras1/2 and the G protein-coupled receptor Gpr1. The subsequent increase in cAMP levels triggers the dissociation of Bcy1 from the Tpks allowing them to phosphorylate their substrates (7, 8). Additionally, cAMP-independent activation of PKA has been reported (9). In addition to cell growth, Tpk effectors influence many other processes including carbohydrate metabolism, cell cycle progression, sporulation, pseudohyphal development and longevity by controlling the activities of metabolic enzymes, transcription and autophagy factors (10–13).

Similarly to TORC1, TORC2 phosphorylates and activates AGC-kinases, including Ypk1 and its redundant paralog Ypk2, in their hydrophobic motif (14, 15). Deletion of *YPK2* produces no obvious phenotype suggesting that it only plays a minor role in cell growth control (16). Recent work has demonstrated that TORC2 is regulated downstream of membrane tension (17), oxidative stress (18) and carbon cues (19). In turn, Ypk1, which is homologous to the mTORC2 substrate SGK in humans, couples TORC2 signals to the regulation of membrane lipid biosynthesis and the regulation of cell surface area (20, 21).

Despite their central role in the regulation of fundamental cellular processes and intense efforts in understanding the functions controlled by these kinases, a systematic assessment of their targets has been lacking to date. A previous study aimed at systematically defining changes in the phospho-proteome associated with the absence of protein kinases, using kinase deletion strains (22). However, this approach is limited to the study of non-essential kinases and allows cells to adapt to the absence of a kinase.

To overcome these limitations, we employed yeast strains expressing analog-sensitive (23) variants of PKA, Sch9 and Ypk1. Mutation of the gatekeeper residue of these kinases allows binding of the bulky ATP-analog C3-1’-naphthyl-methyl PP1 (1NM-PP1), thus preventing ATP-binding and rendering the enzymes inactive. Using this selective and acute way of kinase inhibition, we explored the phospho-protein targets downstream of each of these major AGC-kinases by means of quantitative mass spectrometry. In these phospho-proteomics datasets we identified both known and potentially new targets of each tested kinase. As expected, we found extensive substrate overlap between PKA and Sch9. Unexpected was our finding that several substrates were shared between PKA and/or Sch9 and Ypk1 and that many sites hypo-phosphorylated upon Ypk1 inhibition resided in an RRxS/T-motif, which has previously been associated with PKA and Sch9, rather than Ypk1.

Among the numerous potentially new kinase-substrate relationships discovered in this study, we chose neutral trehalase Nth1 as a candidate for follow-up experiments. Nth1 has been employed as a model PKA-substrate in multiple previous studies (24, 25). Trehalose is a disaccharide that functions as a stress-protectant and reserve carbohydrate under adverse conditions (26, 27). Upon return to favourable conditions, trehalose is converted into two molecules of glucose by trehalases, including the neutral trehalase Nth1 (25, 28). The regulation of Nth1 by PKA has long been recognized as important for cell survival (24, 29). Here, we site-specifically validated that Ypk1-, as well as PKA-inhibition reduced its phosphorylation of an RRxS-site and showed that this is associated with reduced trehalase activity.

These findings highlight the need for revisiting the Ypk1 consensus motif and prompt further investigation of the relationship of PKA and TORC1-Sch9 signalling on one hand and the TORC2-Ypk1 pathway on the other hand.

## RESULTS

### Evaluation of the chemical genetics approach for the analysis of *S. cerevisiae* AGC-kinases

To study substrate specificities and potential signal convergence of growth-regulatory AGC-kinases we used quantitative mass spectrometry to determine changes in the yeast phospho-proteome after acute inhibition of Sch9, PKA or Ypk1. As overlapping functions of PKA and Sch9 have been reported (3, 6), we also performed simultaneous inhibition of PKA+Sch9. Genes encoding *wt* kinases were replaced with alleles encoding the corresponding analog-sensitive enzymes (34). In the *pka^as^* strains all three PKA-paralogs (*TPK1, TPK2, TPK3)* were replaced with analog-sensitive alleles. The *YPK1* paralog *YPK2*, which is considered to play a minor role compared to *YPK1* (16), was deleted in all strains to avoid redundancy. *S. cerevisiae* strains with analog-sensitive Ypk1 (18), Sch9 (30) and PKA (31) have been used successfully in previous studies.

We characterized these analog-sensitive strains (denoted as *sch9^as^*, *pka^as^*, *ypk1^as^*and *pka^as^*+*sch9^as^*) versus the isogenic background (referred to as *wt^as^*) by assessing their growth on plates with and without ATP-analog (Fig. 1 and Fig. S1). Given the central role of all three kinases in growth regulation, we expected visible effects on colony size upon the inhibition of any of them. A slight reduction of colony size was observed for the *sch9^as^* and *pka^as^*+*sch9^as^* strains even on plates without 1NM-PP1, indicating that these strains were mildly hypomorphic. The strongest effect of kinase inhibition was seen in case of the *pka^as^*+*sch9^as^*strain, which did not form colonies on plates containing 500 nM 1NM-PP1. Inhibition of Sch9 alone also exhibited a visible reduction in growth. Interestingly, a small number of regular-sized colonies formed under these conditions, indicating that a limited number of cells adapted or became insensitive to Sch9-inhibition. This is consistent with a previous observation in our lab of an *sch9Δ*-strain rapidly generating adapted clones (32). While less pronounced than for Sch9, the inhibition of Ypk1 also had a strong effect on colony formation on plates.

**Fig. 1:**
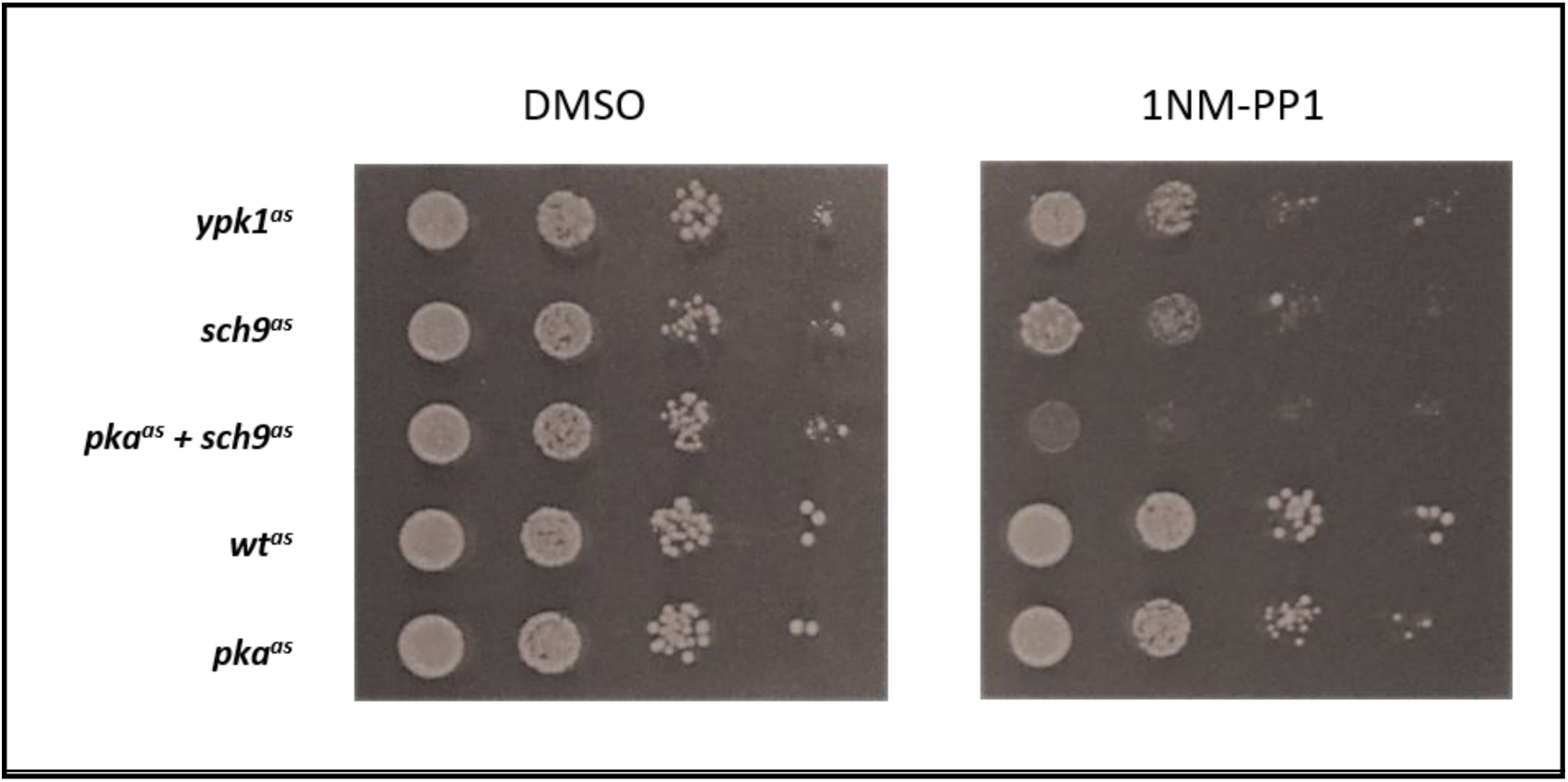
Inhibition of AGC-kinases inhibits colony formation to varying extents. Analog-sensitive strains as indicated were spotted on YPD plates containing DMSO or 500 nM 1NM-PP1 in 10x serial dilutions and imaged after two days at 30°C.

In contrast, the presence of 1NM-PP1 only mildly compromised colony size in case of the *pka^as^* strain. Therefore, the analog-dependent inhibition of *pka^as^* is either less efficient than for *sch9^as^* and *ypk1^as^* or the absence of PKA activity is compensated by other pathways in our strain background. This second alternative seems unlikely in light of the observation that *tpk1Δ tpk2Δ tpk3Δ* triple-deletion strains are not viable (6,8,16). It can be envisioned, however, that there is an increased likelihood of an adaptive response upon prolonged exposure of an analog-sensitive strain to ATP-analog than during creation of a deletion strain. Nevertheless, the comparison of the 1NM-PP1 treated *sch9^as^*and *pka^as^*+*sch9^as^* strain demonstrated that *pka^as^* is sensitive to 1NM-PP1 (Fig.1 and Fig. S1) and this finding further suggested that the compensation for PKA function is mainly through Sch9, which is consistent with previous findings (see Broach, 2012 for a review).

### PKA-, Sch9- and Ypk1-dependent phospho-proteomes

For phospho-proteome analyses, analog-sensitive strains treated with 1NM-PP1 vs. DMSO were compared by label-free, quantitative shotgun proteomics.

At a false-discovery rate (FDR) of 0.01, 7575 peptides, 4628 of which were phosphorylated (61%), were identified from phospho-peptide enriched samples from the respective strains. These phospho-peptides corresponded to 6373 phospho-sites (at site-decoy fraction: 0.01), mapping to 1430 proteins.

Out of the 4628 identified phospho-peptides, 3113 had no more than one missing value in three replicates of any sample and were thus considered for further analysis (Tab. S1). At a multiple-testing corrected p-value cut-off (p_Adj_) of 0.05, 439 phospho-peptides were found to be altered in their abundance upon 1NM-PP1 treatment in the Ypk1-, 128 in the Sch9-, 118 in the PKA- and 293 in the PKA+Sch9 dataset (i.e. 2.5-9.5% of phospho-peptides were altered in the datasets). For comparison, 12 phospho-peptides were classified as changing when *wt^as^* cells were treated with the ATP-analog, corresponding to a 0.4% false-positive rate.

Principal component analysis of these data revealed that replicate samples clustered and that separation of samples was mainly explained by genotype rather than by treatment (Fig. S2). While the *pka^as^* strain was localized near *wt^as^*, the *ypk1^as^* strain on one hand and *sch9^as^* and *pka^as^*+*sch9^as^* strains on the other formed separate clusters.

With respect to the *sch9^as^* and *pka^as^*+*sch9^as^* strains this result is consistent with the observed reduced colony size of these strains obtained in a spot assay in the absence of 1NM-PP1 (Fig. 1 and Fig. S1). This suggests that the gatekeeper mutations in Sch9 and Ypk1 reduce the catalytic activity of these kinases and render the strains hypomorphic. In contrast, we did not observe impaired colony formation of the *ypk1^as^*strain. This indicates that the extent of reduced phosphorylation of Ypk1 targets in this strain is not sufficient to lead to an observable proliferation phenotype.

A heatmap depicting the intensity fold changes of peptides significantly (p_Adj_ < 0.05) hypo- or hyper-phosphorylated upon kinase inhibition is shown in Fig. S3. This global overview suggests a similar pattern of changes in the PKA+Sch9, Sch9 and PKA datasets, albeit for some phospho-peptides changes are only apparent in the PKA+Sch9 dataset. In contrast, the Ypk1 dataset contains some phospho-peptides which change in the same direction as in one or more of the other datasets, but the majority appears to change uniquely in this sample.

To further assess the commonalities and differences between the inhibition of these kinases we determined the overlap among significantly (p_Adj_ < 0.05) altered phospho-peptides observed for each strain tested.

We first asked whether our data would recapitulate the previously reported interaction between PKA and Sch9 signalling in the form of shared targets (3). Indeed, one third to one half of the phospho-peptides significantly affected (i.e. hyper- or hypo-phosphorylated) in one of the datasets was also significant in the other (50 out of 117 and 128 respectively). This may indicate a strong overlap of or cross-talk between the signalling pathways. In the former case it can be envisioned that for some shared targets the inhibition of one of the inputs is compensated by the other. An argument in favour of this proposition is the finding that approximately twice as many significant phospho-peptides (293) were identified in the PKA+Sch9 dataset compared to the individual PKA and Sch9 datasets. Furthermore, 112 of these were unique to the PKA+Sch9 dataset, consistent with the notion that wild type PKA or Sch9 masks the loss of the other for many targets. On the other hand, 80% of the Sch9- and 66% of the PKA-targets were significantly altered also in the PKA+Sch9 dataset, arguing that the high number of sites unique to PKA+Sch9 is not merely due failure to detect true positives repeatedly over datasets.

We next addressed the overlap between PKA/Sch9- and Ypk1-targets, which we expected to be low as TORC2-signalling is considered to be less connected to PKA- and Sch9-signalling than the latter two to each other. Consistent with this idea, we found 297 phospho-peptides uniquely affected by Ypk1 inhibition. Surprisingly, 142 phospho-peptides (32%) were shared with at least one other dataset, especially with PKA+Sch9 (27%).

The results were qualitatively similar when focussing only on hyper- (Fig. S4B) and hypo-phosphorylated (Fig. S4C) phospho-peptides, in that targets unique to the Ypk1- and PKA+Sch9-datasets constituted the largest groups while a strong prevalence of phospho-peptides shared between sets was also apparent.

A full list of gene ontology (GO) terms significantly enriched (p < 0.05) among proteins with regulated phospho-sites is given in Tab. S2. Among the Biological Process GO terms, ’endocytosis’, which was enriched in the Ypk1-dataset, is a major downstream process regulated by this kinase (33). Indeed, the majority of the 65 proteins associated with this term form an interconnected network with Ypk1 that contains key proteins from several stages of endocytosis (initiation: Syp1, Ede1; budding: Pan1, Las17, Ent1, Ent2; and coat disassembly: Inp52) (Fig. S5)(34). It should be noted that many of these proteins likely are not direct Ypk1 targets, but connected further downstream in the Ypk1-Fpk1-Dnf1/Dnf2-Akl1 axis (Fig. S5) (34). Beyond these fundamental endocytic steps, proteins involved in cargo recruitment (Bul1, Ecm21, Rod1, Aly1 and Aly2) were also found to be affected by Ypk1 inhibition. Interestingly, the nutrient permease regulator Npr1 forms part of this network, even though it previously had been associated with TORC1 rather than TORC2 signalling (35).

’Regulation of transcription involved in meiotic cell cycle’ and ‘Response to abiotic stimulus’ were both enriched in the PKA-, while no significant terms were found for the Sch9- and PKA+Sch9-datasets.

Similar gene ontologies related to transferase/glucosyltransferase activity were detected for the PKA, Sch9 and PKA+Sch9 dataset and the proteins associated with these GO-terms in the three datasets strongly overlapped and included subunits of the trehalose 6-phosphate/phosphatase complex (Tsl1 and Tps3), glycogen phosphorylase Gph1 and glycogen synthases Gsy1 and Gsy2 (Tab. S2). The fact that both homologs Gsy1 and Gsy2 were detected based on distinct peptides lends validity to the finding that the regulation of storage carbohydrate metabolism is an important function of PKA and Sch9 signalling (Tab. S1). Additionally, the neutral trehalase Nth1 was found to be affected by PKA and Ypk1 inhibition (see below).

In summary, we observed that all the analog-sensitive kinases were affected by 1NM-PP1 addition based on the growth of the corresponding strains and on observed changes in protein phosphorylation. These phosphorylation changes are consistent with previous knowledge, especially with respect to the close connection between PKA- and Sch9-signalling. Additionally, enriched GO-terms reflect previous data, but are represented also by additional proteins that had not been linked to the respective kinases.

### Sequence motifs

To determine if AGC-kinases in our experiments preferentially targeted specific amino acid sequences, we generated logos of the sequences surrounding phospho-sites significantly affected by 1NM-PP1 treatment. This was done individually for up- and down-regulated sites and for a combination of both for each of the four datasets (Fig. 2).

**Fig. 2:**
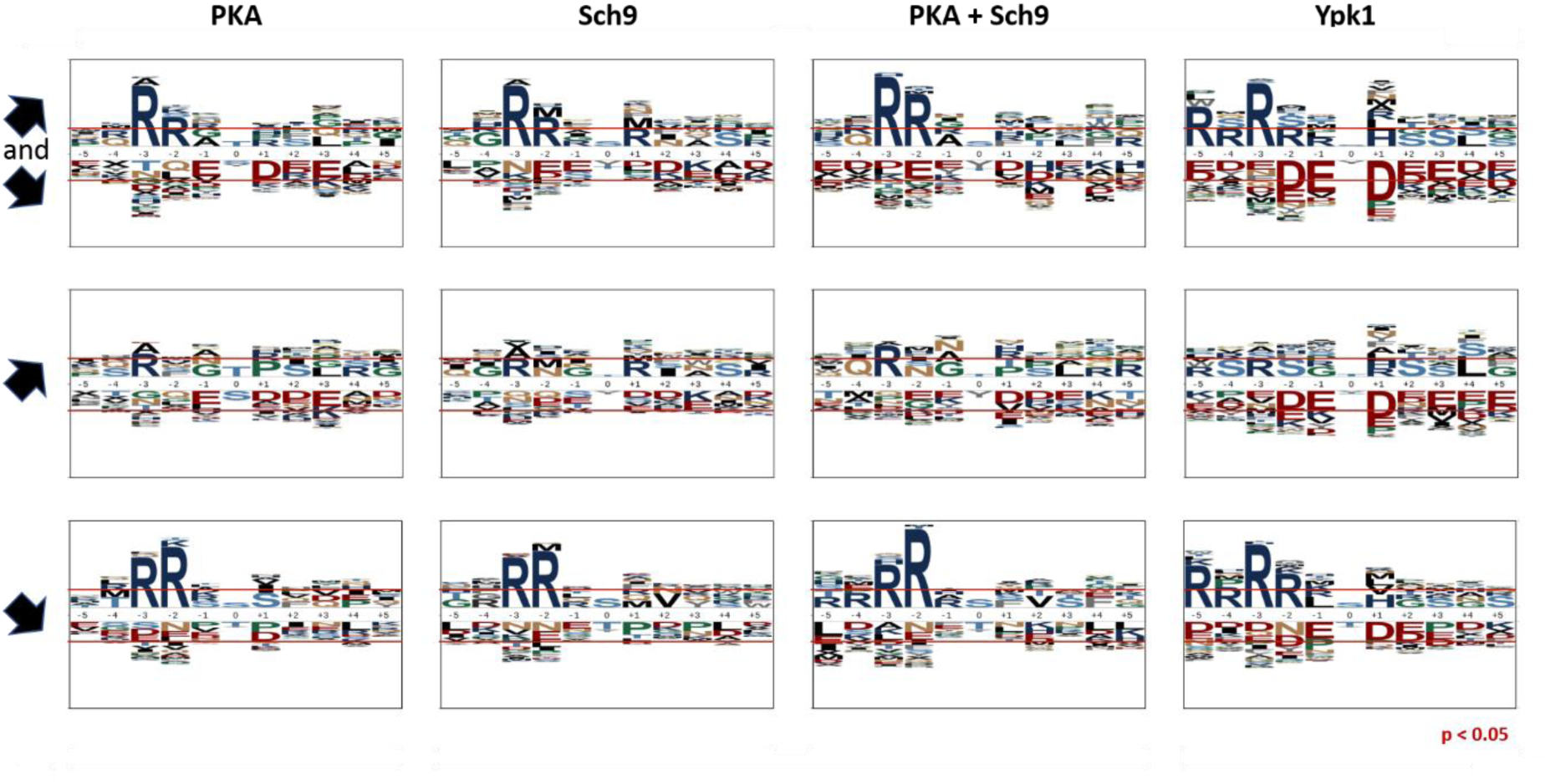
AGC-kinase target sites are located in basic sequence motifs. Sequence motifs from -5 to +5 amino acids surrounding serines and threonines significantly (P_adj_ < 0.05) affected by inhibition of PKA, Sch9, PKA+Sch9 and Ypk1 (left to right). Top to bottom rows: Union of hyper- and hypo-phosphorylated, hyper-phosphorylated, hypo-phosphorylated sites. Red horizontal lines indicate over- and underrepresentation at p = 0.05.

A clear enrichment of arginines in the -2 and -3 position of residues hypo-phosphorylated upon the inhibition of any of the kinases was observed. The motifs of the PKA, Sch9 and PKA+Sch9 datasets closely resembled each other in this respect and exhibited stronger enrichment of arginine in the -2 than -3 position. These findings are consistent with the reported RRxS/T consensus motif for PKA and Sch9 and may therefore reflect the presence of direct targets of the studied kinases (36, 37). For hypo-phosphorylated sites in the Ypk1-dataset in contrast, arginine was more strongly enriched in the -3 position and further enrichment of arginines N-terminal of the phosphorylated residue, in particular at -5 and -2, could be observed. The sequence context determined for Ypk1 fitted the reported RxRxxS/T-motif due to the enrichment of arginine in the -5 and -3 position (38), however additionally an unexpectedly high frequency of arginine in the -4 and especially -2 position was apparent in our data .

It is noteworthy that a slight enrichment of arginine in the -3 position of sites hyper-phosphorylated upon kinase inhibition was observable. As the change in abundance was opposite to what was expected for direct targets, it is possible that this residue forms part of the recognition sequence of a kinase that is itself repressed by the analog-sensitive kinases studied.

In each case, the PKA+Sch9 motifs constituted a more pronounced version of the PKA and Sch9 motifs, therefore indicating that Sch9- and PKA-signalling converge either through shared targets or cross-talk.

### Phosphorylation levels of known direct AGC kinase targets

To estimate if our approach was suitable to identify direct AGC-kinase targets we asked how many of their previously known target-sites were identified and exhibited significant down-regulation upon kinase inhibition in our dataset. Ypk1, Sch9 and PKA target sites derived from the literature are listed in tables S3A-C.

While only 14% to 19% of *bona fide* kinase target sites were detected (i.e. identified and quantified) by mass spectrometry, the vast majority of these sites were significantly hypo-phosphorylated upon inhibition of the respective kinase:

Three of the 21 known Ypk1 target sites were detected in this study and all found to be down-regulated upon Ypk1-inhibition (Lag1 S24, Gpd1 S24, Rod1 S138; Fig. S6A). Similarly, two of the 13 known Sch9 target sites were detected here and both were down-regulated in the Sch9 and the PKA+Sch9 datasets (Stb3 S254, Maf1 S90; Fig. S6B). Finally, six of 31 *bona fide* PKA-sites were detected and three of them were down-regulated in the PKA (Bcy1 S145, Nth1 S60, Nth1 S83) and six in the PKA+Sch9 (Bcy1 S145, Nth1 S60, Nth1 S83, Maf1 S90, Pat1 S456 and Pat1 S457) dataset (Fig. S6C).

Therefore, while our phospho-proteomics data suffered from the high false negative rate with respect to detection, typically associated with this approach, the quantitative accuracy appeared sufficient to detect biologically meaningful changes.

### Many proteins possess more than one regulated phospho-site

Several of the previously reported AGC-kinase targets (e.g. Fpk1, Orm1, Fps1, Stb3, Maf1 and Rim15) harbour more than one regulated phospho-site (Tab. S3A-C) (39–44). We therefore asked if this feature could also be observed in our study. Site-specific quantification of phosphorylation levels can be complicated by several sites residing on the same phospho-peptide as it may not be clear which site is responsible for an observed change in phospho-peptide abundance. Furthermore, sites located near lysine or arginine may impede tryptic cleavage and distort the quantification of nearby residues. We therefore used the presence of arginine in the -3 position from the phosphorylated residue, which is a frequent feature of target sites of the AGC-kinases under investigation, as an additional criterion for selecting proteins with multiple regulated sites (21, 45).

Notable observations included the detection of two regulated RxxS-sites on the previously reported Sch9-target Stb3 in the Sch9 and PKA+Sch9 datasets (42). We also observed two and three regulated phospho-sites on the protein kinase Ksp1 in the Sch9 and PKA+Sch9 datasets respectively, one of which (S624) was also significant in the Ypk1 dataset. Additionally, S884-phosphorylation on this protein was only affected upon Ypk1-inhibition. In the Ypk1 dataset, only a few of the RxxS/T-sites on multi-site proteins were found in the strict RxRxxS/T-motif ascribed to Ypk1 (45). Instead, several of the multi-site proteins were shared with other datasets, including Smy2, Gcs1, Igo2, Ksp1 and Avt1 (Tab. S1).

Similarly, several of the regulated RxxS/T-sites on multi-site proteins in the PKA dataset were found in relaxed motifs, i.e. without a basic residue in the -2 position, but with a proline at +1 (T70 and S96 on Smy2; T161 and T170 on Gcs1; S135, S147 and S161 in Tsl1). Interestingly, four regulated phospho-sites were detected on the trehalose synthase subunit Tsl1 in all four datasets. Finally, three RxxS phospho-sites on Bcy1 were significantly affected in the PKA and PKA+Sch9 datasets. Phospho-peptides containing either S83 or S145 exhibited an ∼45%-75% reduced abundance upon kinase inhibition in both datasets, while phospho-peptides with T129 showed an up to two-fold increase in the PKA dataset.

As Bcy1 is the common regulatory subunit of all three catalytic PKA-subunits (Tpk1, Tpk2 and Tpk3), it is conceivable that these sites form part of a feedback loop regulating PKA activity. Indeed, S145 has previously been reported as a PKA target as part of its negative feedback mechanism (46). The same study identified S145 as the by far major target of PKA on Bcy1, which is consistent with this site being the only RRxS/T-site in the protein. The other sites determined in our study may therefore be targets of other kinases or weaker PKA-targets.

We did not detect any peptides on phospho-diesterase Pde1, which has been reported as another component of cAMP-signalling that is a target PKA (47). Instead, S534 and S631 on Ras GTPase activating protein Ira2 were significantly altered in their phosphorylation state upon PKA inhibition and may therefore constitute a so far unknown feedback mechanism of PKA.

The reason for the clustering of phosphorylation-sites on AGC-kinase targets is not well understood in most cases, but it may allow generating switch-like responses similarly to other kinase targets (48). In any case, the discovery of multiple sites within certain proteins, which are affected by kinase inhibition suggests that the function of these proteins is regulated by said kinase.

### Phospho-sites within kinase consensus motifs

As described above, we observed a strong enrichment of arginine in the -3 position of phospho-peptides significantly hypo-phosphorylated upon inhibition of any of the kinases tested. Indeed, of the 484 identified phospho-sites within a RxxS/T-motif, 117 were significantly (P_adj_ < 0.05) hypo-phosphorylated in at least one of the datasets. Since this is consistent with previously reported kinase consensus motifs (RRxS/T for Sch9 and PKA and RxRxxS/T for Ypk1) (37, 45), we argued that the corresponding sites may constitute potential direct kinase substrates. We asked if these potential direct targets are shared between kinases tested or specific to individual kinases.

Similar to the data on all hypo-phosphorylated sites, the vast majority of RxxS/T-sites in the PKA or Sch9 dataset were also part of the PKA+Sch9 set. Not all of these sites overlapped between the PKA and Sch9 samples, indicating that they may be targets of only one of the two kinases or that inhibition of one kinase targeting a site is compensated by the second kinase (Fig. 3A).

**Fig. 3:**
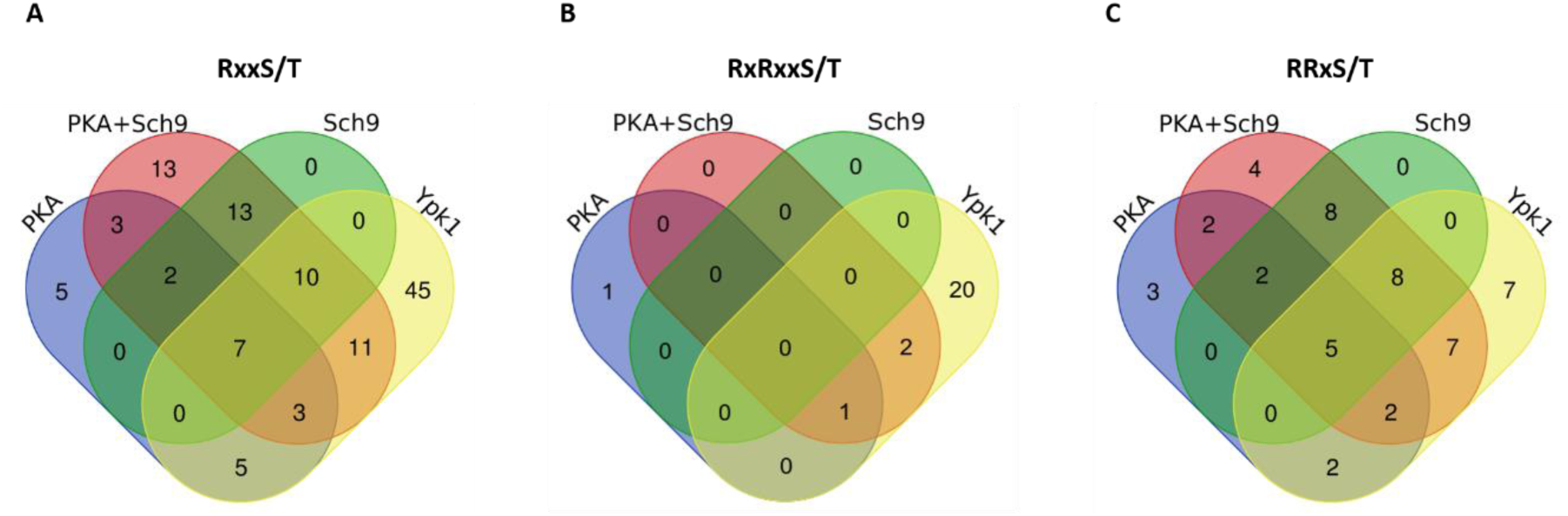
Overlap of hypo-phosphorylated sites associated with kinase consensus motifs between kinase inhibition datasets. Venn-diagrams represent phospho-sites significantly (p_Adj_ < 0.05) hypo-phosphorylated upon inhibition of PKA, PKA + Sch9, Sch9 or Ypk1 in RxxS/T-(A), RxRxxS/T-(B) and RRxS/T-(C) motifs.

While 45 RxxS/T-sites were uniquely affected by Ypk1-inhibition, 36 sites were also hypo-phosphorylated in at least one other dataset. As the Ypk1 consensus motif is considered to be distinct from the PKA/Sch9 motif, we asked if this overlap mainly comprised sites within RRxS/T- or RxRxxS/T-motifs.

We therefore evaluated the overlap of significantly hypo-phosphorylated RxRxxS/T and RRxS/T sites between datasets (Fig. 3B and 3C). As expected, the majority of RxRxxS/T-sites (twenty of 24) were unique to the Ypk1-set. Consequently, sites with an RxRxxS/T-motif do not explain the overlap of RxxS/T-sites between the Ypk1 and the other three datasets.

On the other hand, 31 RRxS/T-sites were hypo-phosphorylated in the Ypk1 dataset, of which 24 overlapped with at least one of the other sets (Fig. 3C). Among these 31 sites only four were simultaneously in an RxRxxS/T-motif (i.e. RxRRxS). Assuming these sites are indeed direct Ypk1 targets, the previously defined Ypk1 consensus motif of RxRxxS/T may need to be reconsidered. We next asked if the RRxS/T-sites that are potential Ypk1 targets had any other sequence characteristics close to the phospho-site, but no obvious amino acid enrichments at other positions within five residues from the phospho-site were observed (Fig. S7).

Following the unanticipated observation that Ypk1 affects the phosphorylation state of sites presumed to be PKA- and/or Sch9-targets based on our results and their sequence context, we next aimed to validate this observation on a selected candidate protein.

### Nth1 phosphorylation of S83 is positively regulated by PKA and Ypk1

Our phospho-proteomics assay detected a phospho-peptide (_81_RGSEDDTYSSSQGNR_95_ + phos) of neutral trehalase Nth1 with potential phosphorylation at S83 or T87 (or a mix of both), down-regulated with a fold-change (FC) of 0.6 (p = 0.007), 0.5 (p = 0.009) and 0.3 (p = 4 × 10^-5^) upon Ypk1, PKA and PKA+Sch9 inhibition respectively. A second phospho-peptide of Nth1 mapping to S60 (_58_TMSVFDNVSPFKK_70_ + phos) exhibited significant down-regulation upon Ypk1 (FC = 0.4; p = 0.0004), PKA (FC = 0.5; p = 0.01) and PKA+Sch9 (FC = 0.5; p = 0.02) inhibition.

Nth1 is a commonly cited model-substrate of PKA and its trehalase activity is increased by PKA-dependent phosphorylation of its N-terminal tail (29,49,50).

In contrast, while Ypk1 had previously been shown to regulate metabolic functions, including of enzymes influencing the levels of the osmo-protective metabolite glycerol, the regulation of the major thermo-protective metabolite trehalose has not been reported (51, 52). To validate our phospho-proteomics results we asked whether Ypk1-inhibition would alter the phosphorylation of an RRxS-site (S83) on Nth1, whose phosphorylation had previously been attributed to PKA (25, 50).

We evaluated the phosphorylation of this site on endogenously 3xFLAG-tagged Nth1 after immuno-affinity purification by western blotting with an antibody against phosphorylated S83 (S83p) (25). Indeed, the signal was reduced upon 1NM-PP1 treatment of strains harbouring either analog-sensitive Ypk1 or PKA (Fig. 4). Importantly, no signal remained in the absence of both enzymatic functions. We also tested the phosphorylation of S83 upon Ypk1, PKA and Ypk1+PKA inhibition in an independent experiment using a targeted proteomics assay. The results closely agreed with the ones obtained by western blotting and confirmed that S83, and not T87, was the phosphorylated residue (Fig. 5; Fig. S8).

**Fig. 4:**
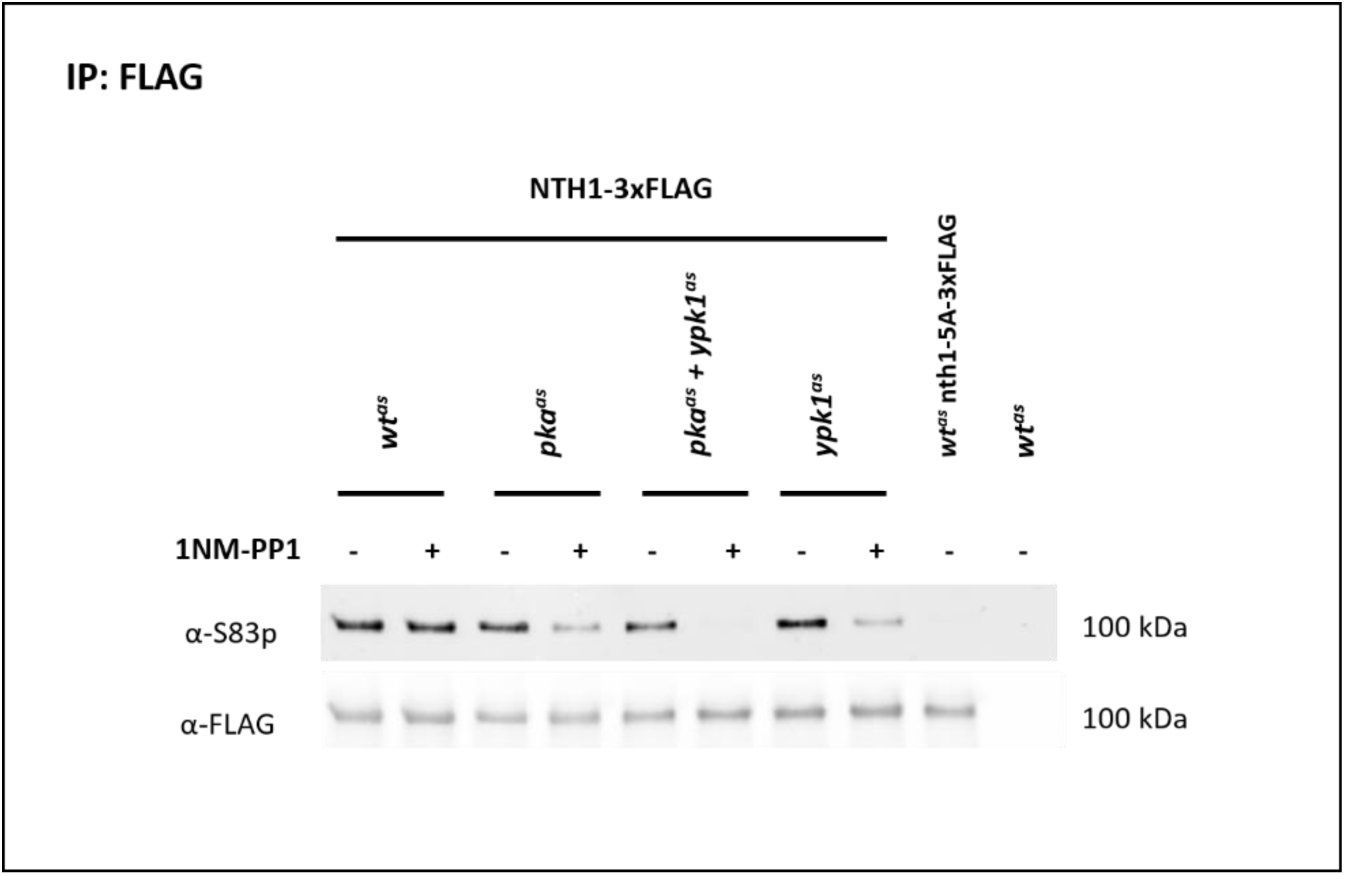
Western blotting demonstrates dephosphorylation of Nth1 S83 upon inhibition of PKA, PKA+Ypk1 and Ypk1. Strains *wt^as^*, *pka^as^*, *pka^as^*+*ypk1^as^* and *ypk1^as^* harbouring endogenously C-terminally 3xFLAG tagged Nth1, the corresponding isogenic wt strain with 3xFLAG-tagged nth1-S20A,S21A,T58A,S60A,S83A (*wt^as^* nth1-5A-3xFLAG) and untagged isogenic wt strain (*wt^as^*) were treated with DMSO or 500 nM 1NM-PP1 for 15 min. Eluates from immuno-precipitates with an anti-FLAG antibody were probed with antibodies against S83p and against the FLAG-tag.

**Fig. 5:**
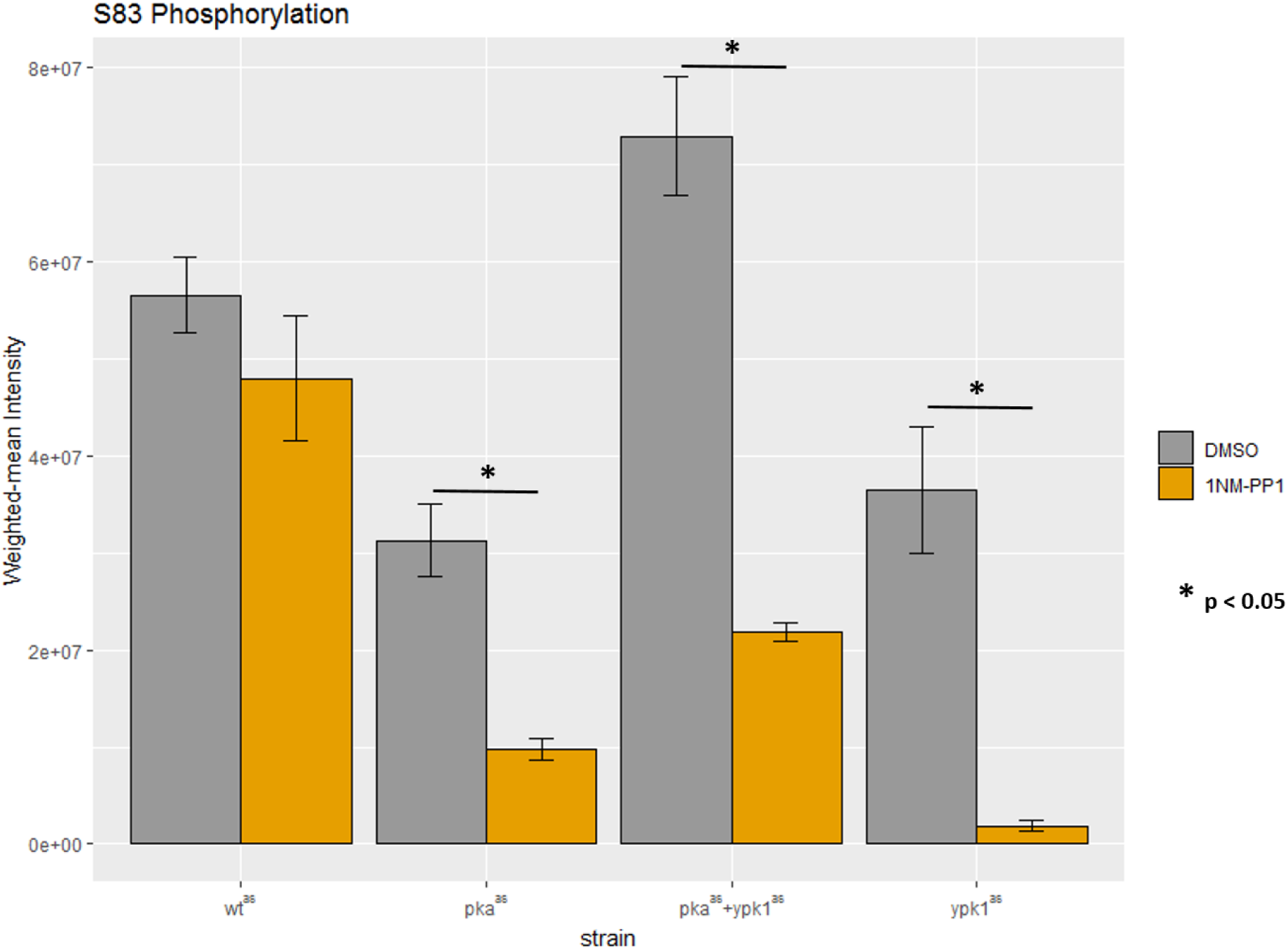
Parallel reaction monitoring demonstrates dephosphorylation of Nth1 S83 upon inhibition of PKA, PKA+Ypk1 and Ypk1. Barplot depicting weighted-mean intensities of phospho-peptides corresponding to Nth1 S83p. Indicated strains were treated with DMSO (grey) or 500 nM 1NM-PP1 (orange) for 15 min. Bars represent averages from three replicates and errorbars represent standard error of the mean. Asterisks indicate significant fold changes at p-values less than 0.05.

These findings validate the effect of Ypk1 on phosphorylation of a site located within an RRxS-motif and therefore not assumed to be a direct Ypk1 target based on its previously reported consensus motif (45).

### Nth1 is activated in a PKA- and Ypk1-dependent manner

It has been previously reported that the PKA-dependent phosphorylation of the N-terminal tail of Nth1 stimulates the enzymès trehalase activity (24,25,29). Based on our results of Ypk1-dependent phosphorylation of Nth1 we asked if Ypk1 would also regulate Nth1 activity. For this we again compared *wt^as^*, *pka^as^* and *ypk1^as^* strains treated for 15 min with 1NM-PP1 with mock-treated controls and assessed trehalase activity by a method similar to the one described by De Virgilio, 1991 (53).

We employed a *nth1Δ* strain as a control in this experiment. For this strain no detectable trehalase activity could be observed (Fig. S9A and S9B) indicating that Nth1 is the only trehalase active under these conditions. Note that acidic trehalase is not expected to be active at the neutral pH used for this assay. We observed that trehalase activity was strongly reduced in the *pka^as^*, and somewhat also in the *ypk1^as^*strain, even in the absence of 1NM-PP1, indicating the hypomorphic nature of these strains (Fig. S9A and S9B). Importantly, a significant reduction of trehalase activity in both analog-sensitive strains was observed upon 1NM-PP1 addition relative to mock treated controls (p = 6 × 10^-6^ and 1 × 10^-5^ respectively; Fig. 6, Fig. S9A and S9B). This indicates that, as for PKA, Nth1-activity was positively regulated in a Ypk1-dependent manner.

**Fig. 6:**
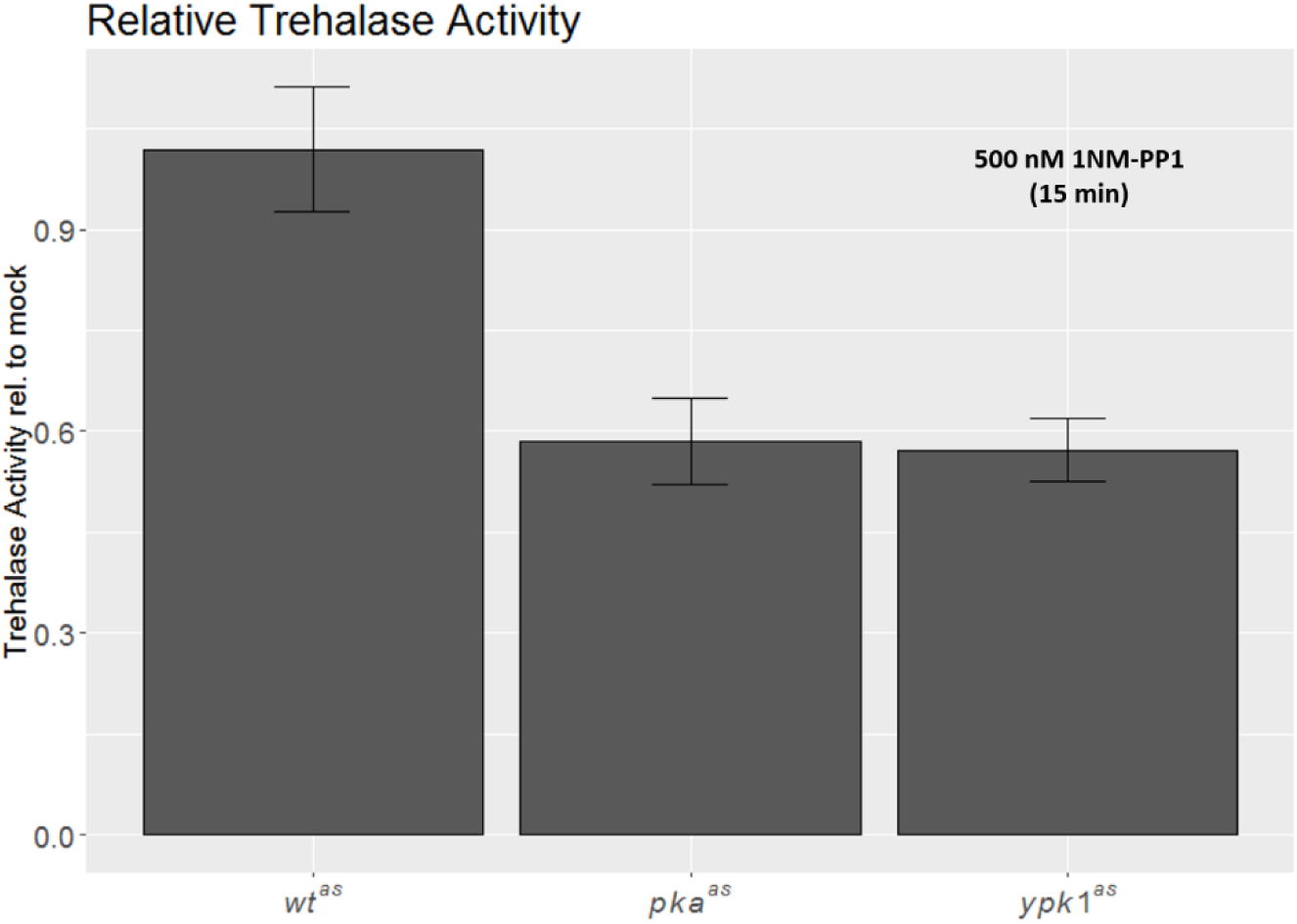
Inhibition of PKA and Ypk1 leads to reduced Nth1 activity. Barplot depicting trehalse activity of 1NM-PP1 treated *wt^as^*, *pka^as^*and *ypk^as^* strains relative to activity of corresponding DMSO-treated strains. Strains were in exponential growth phase and treatment was for 15 minutes. Error-bars depict standard deviation of six replicates. Reduction of relative trehalase acitivity in *pka^as^* and *ypk1^as^*strains is significant with p = 6 × 10^-6^ and p = 1 × 10^-5^ respectively.

It can therefore be envisioned that stresses inactivating Ypk1 cause trehalose accumulation. Similarly to PKA, return to favourable growth conditions and re-activation of Ypk1 may lead to trehalose degradation by Nth1.

## DISCUSSION

### Previous studies on AGC-kinase functions

In this study we investigated the targets of three major growth-regulatory AGC-kinases in *S. cerevisiae*, PKA (Tpk1, Tpk2, Tpk3), Sch9 and Ypk1. For this purpose we employed analog-sensitive versions of these kinases in combination with phospho-proteomics. A *ypk1Δ ypk2Δ* double- and *tpk1Δ tpk2Δ tpk3Δ* triple-deletion renders cells inviable and deletion of *SCH9* leads to severe phenotypic effects, demonstrating the important roles that these kinases perform (6,8,16). A previous large scale phospho-proteomics investigation of strains in which individual kinases had been disrupted included *tpk1Δ, tpk2Δ, tpk3Δ, ypk1Δ* and *ypk2Δ* strains (22). Of necessity, this approach precluded the study of strains harbouring deletions of essential paralogs. A further disadvantage of using deletion strains for studying an enzyme is that such strains inevitably adapt in ways that often mask effects on direct targets.

Analog-sensitive strains of AGC-kinases have been described previously, but not to determine the effect of their inhibition on global phosphorylation changes (17,30,54). We reasoned that the acute inhibition of kinases via a chemical genetics strategy in combination with a direct readout of phosphorylation changes should enable us to increase our understanding of signalling downstream of these kinases and in particular identify novel direct targets.

### Interplay between Sch9- and PKA-signalling

There is ample genetic evidence for a close link between PKA- and Sch9-dependent signalling (3): In fact, Sch9 was identified as a high copy suppressor of a temperature sensitive mutation in the Ras2-guanine nucleotide exchange factor Cdc25 (6). On the flip-side, Sch9 exhibits synthetic lethality with components of the PKA-pathway, Gpr1, Gpa2 and Ras2 (55–57). Signalling through the PKA-pathway represses stress-protective activities of Yak1 and Msn2/4 and G0-entry via Rim15. The subsequent heat-shock sensitivity of an activating mutation in *GPA2* is suppressed by deletion of *SCH9* (58).

Consistently, we observed a strong overlap of phospho-sites affected by PKA- and Sch9-inhibition. Further, a high number of sites detected only upon simultaneous inhibition of both kinases may comprise further shared sites for which the activity of one kinase masks the inhibition of the other. The consensus sequence detected for both kinases closely resembles their established RRxS/T-motif (37,59,60).

These observations pose fundamental questions for future research: Is the considerable overlap between sites affected by PKA and Sch9 inhibition due to cross-talk between the two signalling pathways or due to shared targets of the two kinases? In case of targets unique to one of the kinases, how is this selectivity established, considering the high similarity of the PKA- and Sch9-consensus motifs? The data provided in this study serves as a starting point to answer these questions.

### Interplay between Ypk1- and PKA/Sch9-signalling

While evidence for an overlap between PKA- and TORC1-Sch9-signalling had been reported, TORC2-Ypk1-signalling is considered to be distinct from the former two pathways. Despite shared subunits between the TOR-complexes, TORC1 is believed to mainly respond to nutritional cues while TORC2 presumably receives its input primarily from the cell membrane (61).

Several findings in this study tie Ypk1 much more closely to PKA and Sch9 than we previously assumed: while many phospho-peptides were affected uniquely upon Ypk1 inhibition, a surprisingly high number overlapped with the other datasets. Secondly, while the Ypk1 consensus motif derived in this study was consistent with the previously reported RxRxxS/T-motif, a clear enrichment of arginine in the -2 position could be observed. Closer investigation of basophilic sites hypo-phosphorylated upon Ypk1 inhibition revealed two nearly mutually exclusive sets of Ypk1 targets: one set of sites associated with the RxRxxS/T-motif and almost uniquely affected by Ypk1-inhibition and the other of sites associated with the RRxS/T-motif and largely shared with at least one other kinase. These observations suggest that Ypk1 recognizes two related, but distinct motifs. Alternatively, Ypk1 may regulate these sites indirectly through an RRxS/T-specific kinase.

### Kinase target motifs

The commonly cited Ypk1 target motif is RxRxxS/T (with an additional preference for a hydrophobic residue in the +1 position) and was defined by the Thorner lab based on nine *bona fide* Ypk1 target sites on five proteins and previously published *in vitro* assays on synthetic peptides (46). This resembles the consensus motif of the presumed human homolog of Ypk1/2, SGK (21). It is interesting to note that, while one of the *in vitro* studies to define the Ypk1 target motif did not test peptides containing the RRxS/T-sequence (21), the second reported a Ypk1 motif similar to our study: while arginine in the -3 position was most strongly enriched, a similar prevalence of arginine in -5 and -2 was also observed (37). Parenthetically, for Ypk2 the target sequence even more closely resembled an RRxS/T-motif. Collectively, these results suggest that Ypk1 also recognizes RRxS/T-motifs besides RxRxxS/T.

After the initial definition of the Ypk1 consensus motif (38) twelve further sites on six proteins were identified, which all reside in RxRxxS/T-motifs (38,38,41,62). However, the presence of this motif in each case constituted a criterion in the search for new Ypk1 sites, biasing the selection. It should also be noted that the assays performed in our study targeted Ypk1 during conditions of basal activity rather than in contexts where it is highly activated. It may be speculated that full activation of Ypk1 leads to a change in its substrate preference. This idea is particularly intriguing as distinct phosphorylation steps by Pkh1/2 and TORC2 may lead to partial and full activation of Ypk1 (14, 21).

In contrast to Ypk1, the sequence motif for sites hypo-phosphorylated upon PKA-inhibition closely agrees with the previously reported PKA-consensus motif. Similarly, our Sch9-motif is in agreement with a motif previously defined by peptide-array (37) and with the fact that *bona fide* Sch9 target sites are generally in an RRxS/T-motif (2,32,42).

### Regulation of trehalose metabolism by Ypk1

We further investigated the connection between PKA- and Ypk1-signalling by choosing neutral trehalase Nth1 as an example. The phosphorylation of its N-terminal tail (S20, S21, S60 and S83) by PKA has been previously reported (25,29,50). It has been suggested that this phosphorylation leads to the activation of its trehalase activity, mainly through the binding of 14-3-3 proteins to phosphorylated S60 and S83 (49, 50). We here showed that Nth1 phosphorylation and activity is also diminished by Ypk1 inhibition *in vivo*, therefore establishing a connection betweenTORC2-signalling and storage carbohydrate metabolism.

Only few Ypk1 targets are currently known. Regulation of trehalose metabolism may therefore be a so far overlooked TORC2-dependent stress response. If this regulation is indeed via direct phosphorylation of the N-terminal tail of Nth1 by Ypk1, remains to be determined. The fact that two of the presumed regulatory sites, S20 and S60, reside within an RxRxxS/T-sequence, the preferred Ypk1 target motif, argues in favour of this possibility, regardless of if Ypk1 can also phosphorylate RRxS/T-sites.

Further phospho-sites on trehalose-6-P synthase/phosphatase subunits Tsl1, Tps2 and Tps3 were affected by Ypk1 inhibition and may provide further ways of TORC2-signalling modulating this metabolic pathway.

The TORC2-dependent regulation of endocytosis in contrast is well-established, however mechanistically not well understood (33). Our data provide an extensive list of Ypk1-regulated phospho-proteins involved at several steps of endocytosis. It will be interesting to map the location of these proteins in the TORC2-Ypk1 pathway and to define phenotypic consequences of site-specific mutants of the corresponding phospho-sites.

Finally, we reported phospho-sites affected by the inhibition of AGC-kinases on proteins that had not been previously linked to signalling through these kinases. Based on the close match of sequence motifs derived from these sites and the reported consensus motifs of the kinases it is likely that we detected several direct kinase targets, setting the stage for future focused studies.

## METHODS

### Yeast cultures and assays

*Saccharomyces cerevisiae* strains used in this study are listed in Tab. S4. Strains were constructed using standard yeast genetic manipulation (63).

#### Label-free phospho-proteomics

Three replicates each of DMSO- and 1NM-PP1-(Merck, Darmstadt, Germany) treated *sch9^as^*, *pka^as^*, *pka^as^*+*sch9^as^* and *ypk1^as^* cultures were prepared for label-free phospho-proteomics as previously described (32). In brief, all cultures were diluted to OD 0.2 and grown to exponential phase (OD 0.7-0.8). Each strain was treated with 500 nM 1NM-PP1 or DMSO for 15 minutes after which ice cold 100% TCA was added to a final concentration of 6%. The samples were incubated on ice for 30 minutes. After centrifugation the cells were washed twice with acetone and kept at -70°C. Cell pellets were suspended in a buffer consisting of 8 M urea, 50 mM ammonium bicarbonate, and 5 mM EDTA, then lysed by beat beating. For each biological replicate, 3 mg protein were reduced using 5 mM TCEP, alkylated in 10 mM iodoacetamide, and digested overnight with trypsin (1:125 w/w). Reverse phase chromatography was used to purify samples before phospho-peptide enrichment, which was performed with titanium dioxide resin (1.25 mg resin for each sample) (22). Isolated phospho-peptides were analyzed on an LTQ-Obritrap XL mass spectrometer (Thermo Scientific, Basel, Switzerland). A 90 minute gradient, starting with 3% and ending with 23% acetonitrile, was used for liquid chromatography elution. The 4 most intense ions detected in each MS1 measurement were selected for MS2 fragmentation.

### Data analysis

The 30 .raw files were processed with the MaxQuant software, which uses the Andomeda MS/MS spectrum search engine (64, 65). The following search settings were used: enzyme was Trypsin/P with 2 allowed missed cleavages. The first search tolerance was set to 20 ppm and the main search tolerance to 4.5 ppm. Phosphorylation of serine, threonine and tyrosine residues, oxidation of methionine, acetylation of protein N-termini and deamidation of asparagine and glutamine were used as variable modifications (maximal 5 modifications per peptide). Carbamidomethylation of cysteine was used as fixed modification. The SGD yeast protein database (https://downloads.yeastgenome.org/sequence/S288C_reference/orf_protein/orf_trans_all.fasta.gz) from 2015-01-13 was used to match the spectra. Finally, a peptide and modification site FDR of 0.01 was applied to filter peptide spectrum matches (PSM). Contaminants and decoys were subsequently removed from the results.

The MaxQuant result tables (msms.txt for PSMs and positioning information, modificationSpecificPeptides.txt for intensity values) were imported into the R statistical environment to perform missing value imputation and intensity normalization. Only peptides containing at least one phosphorylation were retained, together with all possible positions of the phosphorylation and their respective scores. All intensity values were log_2_ transformed.

Fig. S10A shows the occurrence of missing values across the 3000+ phospho-peptides revealing that certain peptides and certain MS-runs are more prone to missing values. An example of a peptide with missing values can be seen in Fig. S10B, where all Sch9/PKA-1NM-PP1 values of the peptide HSS(ph)PDPYGINDKFFDLEK and one Sch9 and one PKA value are missing. In the sibling peptide (see below) HSS(ph)PDPYGINDK with clearly higher MS1 intensities all values are present (Fig. S10C). This shows that even if all 3 replicates values are missing, one cannot conclude absence of the corresponding peptide, but these missing values are most likely artefacts inherent to MS1 based quantitation. Phospho-peptides with no missing values in 3 replicates have stronger average MS1 signal intensities compared to peptides with missing values (Fig. S10D). If only one value out of 3 replicate values is missing, the 2 remaining positive replicate values can be used to estimate or impute the missing values (see below). However, if 2 or 3 values are missing, imputation becomes more error prone and can easily lead to false positives, when searching for peptides of differential abundance. Therefore, we only retained peptides with maximally one missing value per 3 replicates. 67.3% of all phospho-peptides fall in this category (Fig. S10E).

We imputed a missing log2-transformed intensity value *IP,S,i* of peptide *P* in replicate *i* of sample *S* by

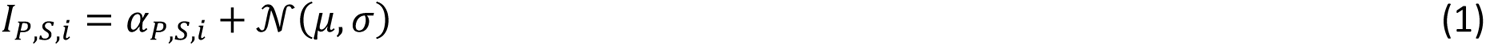

where *α_P,S,i_* is the mean of the 2 non-missing values of peptide *P* in sample *S* and 𝒩(μ, σ) is a Gaussian error term with mean *μ* and standard deviation *σ*. In order to estimate this error term, we first need the notion of sibling peptides. Two peptides are called siblings if the sequence of one peptide is a N/C-terminal extension of the sequence of the other one (i.e. they differ by a missed cleavage) and if they carry the same modifications. It is unlikely that the sample condition has a direct influence on the trypsin cleavage and it can be expected that two sibling peptides have similar expression profiles.

The imputation parameters *μ* and *σ* are then estimated using the sibling peptides present in the MaxQuant results: for each peptide *P* with a missing value in replicate *i* in sample *S*, we searched for sibling peptides *P’* of *P* with no missing values in the 3 replicates of sample *S.* Then we calculated the difference between the *I_P’,S,i_* and *α_P’,S,i_* and used all these differences to estimate *μ* and *σ.* The result shown in Fig. S10F reveals that the missing values are on average slightly, but significantly lower than the average of the 2 non-missing values (t-test p-value of 0.003), a small bias that we took into account in the imputation Equation 1).

After missing value imputation, the R package limma (66) was used for intensity normalization. First the *normalizeBetweenArrays* R-function was used to equalize the log_2_-intensity distributions in all 30 LC-MS runs. The lmFit function was used to remove eventual batch effects present in the 10 samples. The empirical Bayes (eBayes) method was the applied to compare the 1NM-PP1 treated to the control (DMSO) samples for each strain and to calculate fold-changes and p-values adjusted for multiple testing (FDR adjustment).

#### Generation of sequence logos

For each phospho-peptide that exhibited a change upon treatment with an adjusted p-value of less than 0.05 the sequence from -5 to +5 amino acids from each of the phospho-sites localized on the peptide was extracted. In case of ambiguity between isoforms with different phospo-localizations, the peptide with the best localization score was chosen. If a phospho-site was represented by several peptides, the sequence surrounding the site was listed only once. This was performed individually for each strain for hyper- and hypo-phosphorylated peptides and the union of both. Sequence logos were generated with pLogo using all quantified phospho-peptides as a background (67).

#### Gene ontology analysis

For GO-analysis the systematic gene names of proteins harbouring phospho-sites significantly changing upon kinase inhibition with an adjusted p-value below 0.05 were used as a foreground and harbouring any quantified phospho-site as a background. The analysis was performed using the ‘GO Term Finder’ tool at the Saccharomyces Genome Database at a p-value of 0.05 with FDR calculation enabled (68).

The network of proteins associated with GO-term “Endocytosis” in the Ypk1-dataset was displayed the STRING-database tool at default parameters (69).

### Parallel Reaction Monitoring

Samples for targeted proteomics by Parallel Reaction Monitoring (PRM) were prepared from 250 µg of protein in a similar manner as for label-free phospho-proteomics, with the exception that a mixture of magnetic TiO_2_, ZrO_2_ and Ti(IV)-IMAC beads (15 µl each; ReSyn Biosciences, Edenvale, South Africa) were used for phospho-peptide enrichment, essentially following the manufacturer’s instructions.

Data acquisition was performed on an Orbitrap Fusion Tribrid mass spectrometer coupled to an nLC1000 uHPLC. Peptides were separated on a 25 cm EASY-spray column (Thermo Scientific) with a 30 min gradient from 3 to 25% acetonitrile, 0.1% formic acid vs. 0.1% formic acid in water. MS-scans were acquired at 60 000 resolution, with an ion target of 4 × 10^5^ and maximum injection time of 118 ms. Targeted MS/MS employing higher energy collisional dissociation on 9 selected precursors representing Nth1 phospho-peptides and targeted single ion monitoring on two precursors were each acquired at a resolution of 60 000 with an ion target of 4 × 10^5^ and maximum injection time of 200 ms.

For data analysis in Skyline 4.2.0.19072 we employed spectral libraries from previously generated data-dependent acquisition and filtered for seven b- or y-ions from ion 3 to last ion (70). The extracted total areas were normalized to the total ion current area of the corresponding sample and an intensity-weighted mean was calculated for cases in which phospho-sites were represented by several precursors. The significance of changes between DMSO- and 1NM-PP1 treated samples was evaluated using a two-sided Student’s t-test.

### Spot assays

Yeast strains were inoculated in YPD (pH 5.6) and incubated at 30°C and 120 rpm overnight, diluted to OD = 0.1 the next morning and again to 0.0002 in the evening with incubation under the same conditions. Once an OD of about 0.8 was reached cultures were diluted to OD 0.08 and further in a 10x dilution series in the same medium.

For spot assays 3 μl of these cultures were spotted onto YPD agar plates containing 500 nM 1NM-PP1 or an equal volume of DMSO and incubated at 30°C for two days.

### Western blotting

To site-specifically validate changes in Nth1 phosphorylation by western blotting, yeast cultures were treated with 500 nM 1NM-PP1 or the corresponding volume of DMSO for 15 min and harvested by TCA-precipitation as described for phospho-proteomics sample preparation. Yeast pellets were lysed by bead beating (5x 45s at 6500 rpm) in 25 mM Tris buffer containing 6 M urea and 1% SDS. Cleared lysate corresponding to 1.3 mg protein was diluted 1:10 in IP-buffer (50 mM HEPES pH 7.4, 150 mM KCl, 10% glycerol containing protease-(Roche, Basel, Switzerland) and phosphatase-(Thermo Scientific) inhibitors and mixed with 25 µl Epoxy-beads (Thermo Scientific) that had been coupled to anti-FLAG antibody (1.5 µl; Merck) in 0.1 M sodium phosphate buffer containing 1 M ammonium sulfate at 37°C overnight, and washed four times with IP-buffer. The lysate was incubated with the beads at 4°C for 4 h, then washed four times with IP-buffer. Proteins were eluted in 15 µl Laemmli-buffer at 65°C for 10 min, then in further 15 µl Laemmli-buffer at 95°C for 5 min.

The eluates from both steps were combined and resolved on a 7.5% SDS-PAGE gel. Proteins were transferred to a nitrocellulose membrane using an iBlot2 system, blocked in 5% BSA in PBS for 1 h. The membrane was incubated with primary antibodies (1:1000 mouse anti-FLAG (Merck) and 1:100 rabbit anti-Nth1-S21p or anti-Nth1-S83p, kind gifts from the Thevelein lab (25)) in 5% BSA in PBS + 0.1% Tween-20 at 4°C overnight. It was washed four times 5 min in PBS + 0.1% Tween-20, incubated with fluorescently labelled secondary antibodies (LI-COR, Lincoln, Nebraska, USA) in 5% BSA in PBS + 0.1% Tween-20 for 30 min and washed as above. Fluorescent signal was imaged on a LI-COR Odyssey scanner.

### Trehalase activity assays

Measurement of trehalase specific activity was performed as previously described with modifications (53). Six replicates (separate starter cultures) of strains *wt^as^*, *pka^as^*and *ypk1^as^*, as well as one sample of an *nth1Δ*-strain as a control, were inoculated in YPD (pH 5.6) and incubated at 30°C overnight. Cultures were diluted to an OD of 0.05 and incubated for about 10 h before further dilution to OD 0.001 in YPD. The cultures were grown to an OD of approximately 0.7 and, where necessary, cultures were diluted during this time to arrive at a similar OD for all cultures. Subsequently, 100 ml of each culture was centrifuged for 10 min at 2000 g, pellets resuspended in 10 ml YPD and distributed to two 5 ml cultures in 50 ml Erlenmeyer flasks. The cultures were left to recover at 30°C with shaking at 120 rpm for 25 min before treatment with 500 nM 1NM-PP1 (from a 250 μM stock in DMSO) or the corresponding volume of DMSO. After 15 min of shaking at 30°C 0.5 ml of each culture were harvested into 0.5 ml of 0.2 M tricine (pH 7.0) with 0.1% Triton- × 100 and 1x phosphatase inhibitor, 200 nM Okadaic acid (Cell Signaling Technology, Danvers, Massachusetts, USA) and 1 EDTA-free protease inhibitor tablet (Roche) per 30 ml, immediately frozen in liquid nitrogen and stored at -80°C.

Samples were thawed at 30°C in a thermomixer at 1400 rpm, centrifuged for 1 min at 12 000 g and the pellet washed in 1 ml of cold 0.2 M tricine pH 7.0 (1 min, 12 000 g, 4°C). Pellets were resuspended in 1 ml of cold 0.2 M tricine (pH 7.0) and transferred into the wells of a deep-well 96-well plate on ice. The plate was spun for 5 min at maximum speed and 4°C, supernatants removed and pellets resuspended in 320 μl 62.5 mM tricine pH 7.0, 250 μM CaCl_2_ containing protease- and phosphatase-inhibitors. A well which did not contain a yeast pellet was used as blank. The plate and 0.5 M trehalose were equilibrated to 30°C for 5 min before 80 μl of 0.5 M trehalose were added to each sample and incubation at 30°C. After 10, 20, 30 and 40 min a 25 μl aliquot of each sample was transferred into PCR-stripes and heated to 98°C for at least 3 min.

The samples were cooled to room temperature and cell debris spun down briefly. The amount of glucose in the supernatants was determined using a Glucose-Oxidase/Peroxidase kit (Merck) using 5 μl input in a 150 μl reaction volume.

The amount of glucose generated (= 2x amount of trehalose degraded) was determined against a glucose standard and the blank value subtracted from each sample. The slope of the curve of trehalose degraded vs. time after trehalose addition with y-intercept of zero was determined as a measure for trehalase activity.

This activity was normalized to the protein level determined by BCA assay (Thermo Scientific) after spinning the remaining samples and bead beating of the resulting pellets in 400 μl 1% SDS, 6 M urea in 25 mM Tris (pH 6.8). The trehalase activity per protein content of the *pka^as^*and *ypk1^as^* strains were each compared to that of the *wt^as^*strain using a two-sided, unpaired Student’s t-test.

## Acknowledgements

We thank Claudio de Virgilio (University of Fribourg) and Marko Kaksonen (University of Geneva) for helpful advice, Clelia Bourgoint (University of Geneva) for critical reading of the manuscript and Johan Thevelein (KU Leuven) for helpful advice and reagents. The Loewith lab receives funds from the Canton of Geneva, the Swiss National Science Foundation and the European Research Council and is part of the National Center for Excellence in Research for Chemical Biology. We also acknowledge support from PhosphonetX and SignalX group grants of the Swiss National Science Foundation SystemsX programme.

## Supplementary Figures

**Fig. S1:**
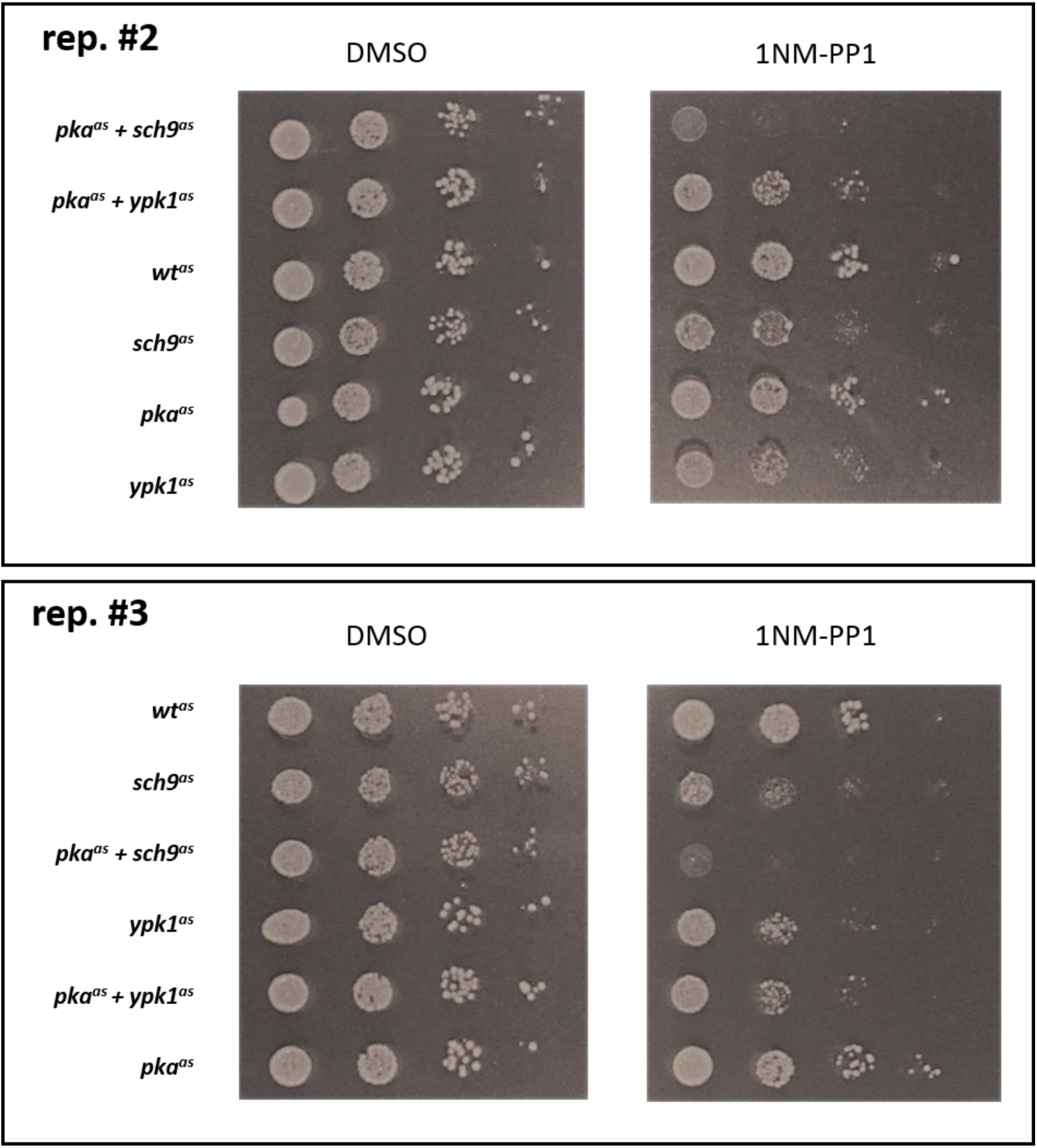
Additional replicates of experiment depicted in Fig. 1. Replicate cultures were processed in parallel to samples shown in Fig. 1A and the double analog-sensitive *pka^as^+ypk1^as^*used in Fig. 5 and S8 was tested in addition.

**Fig. S2:**
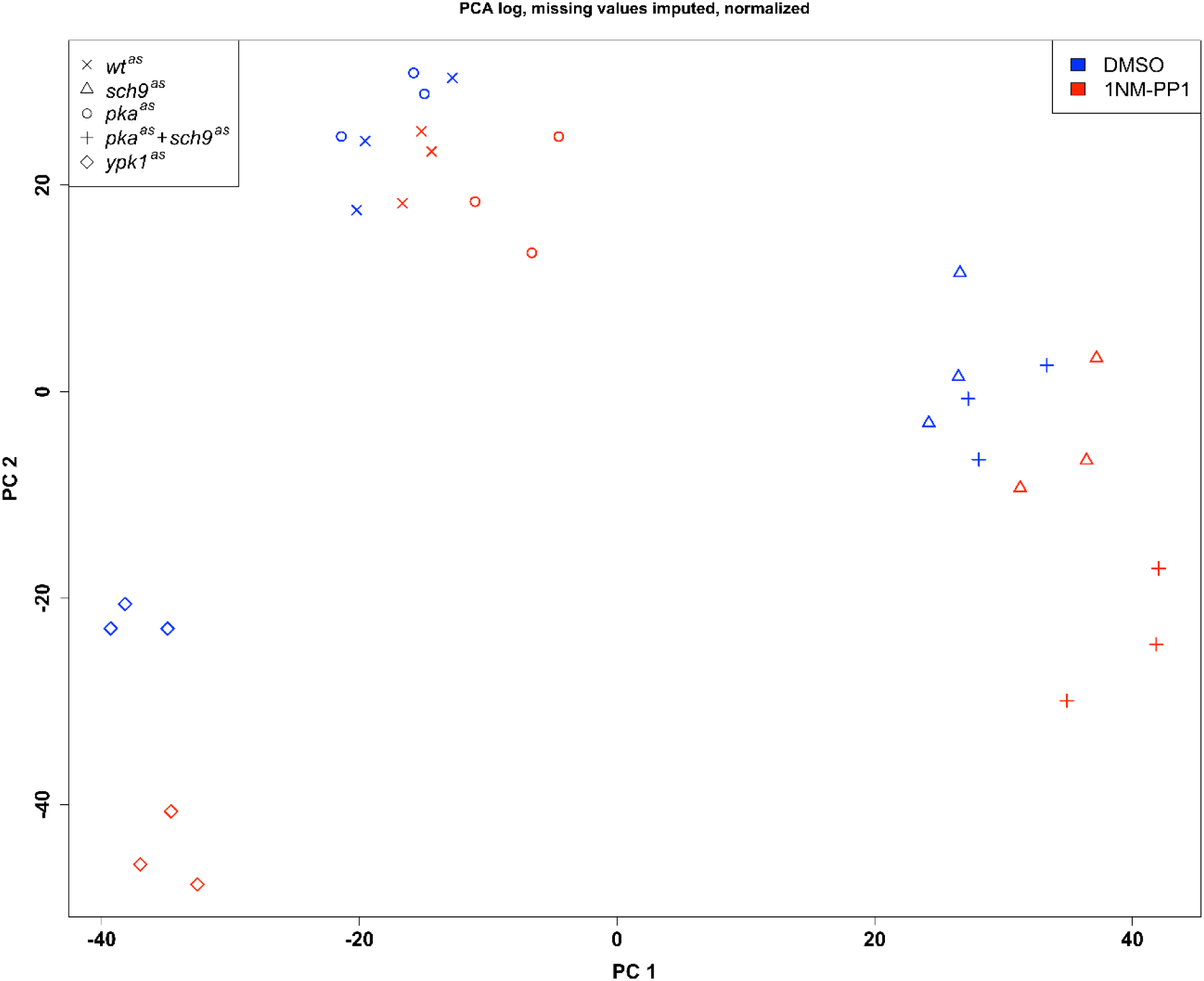
Gatekeeper mutations affect the activity of Ypk1 and Sch9. Components 1 and 2 (PC1 and PC2) of principal component analysis of phospho-proteomics data of *wt^as^* and analog-sensitive strains as indicated after 15 min treatment with DMSO (blue) or 500 nM 1NM-PP1 (red).

**Fig. S3:**
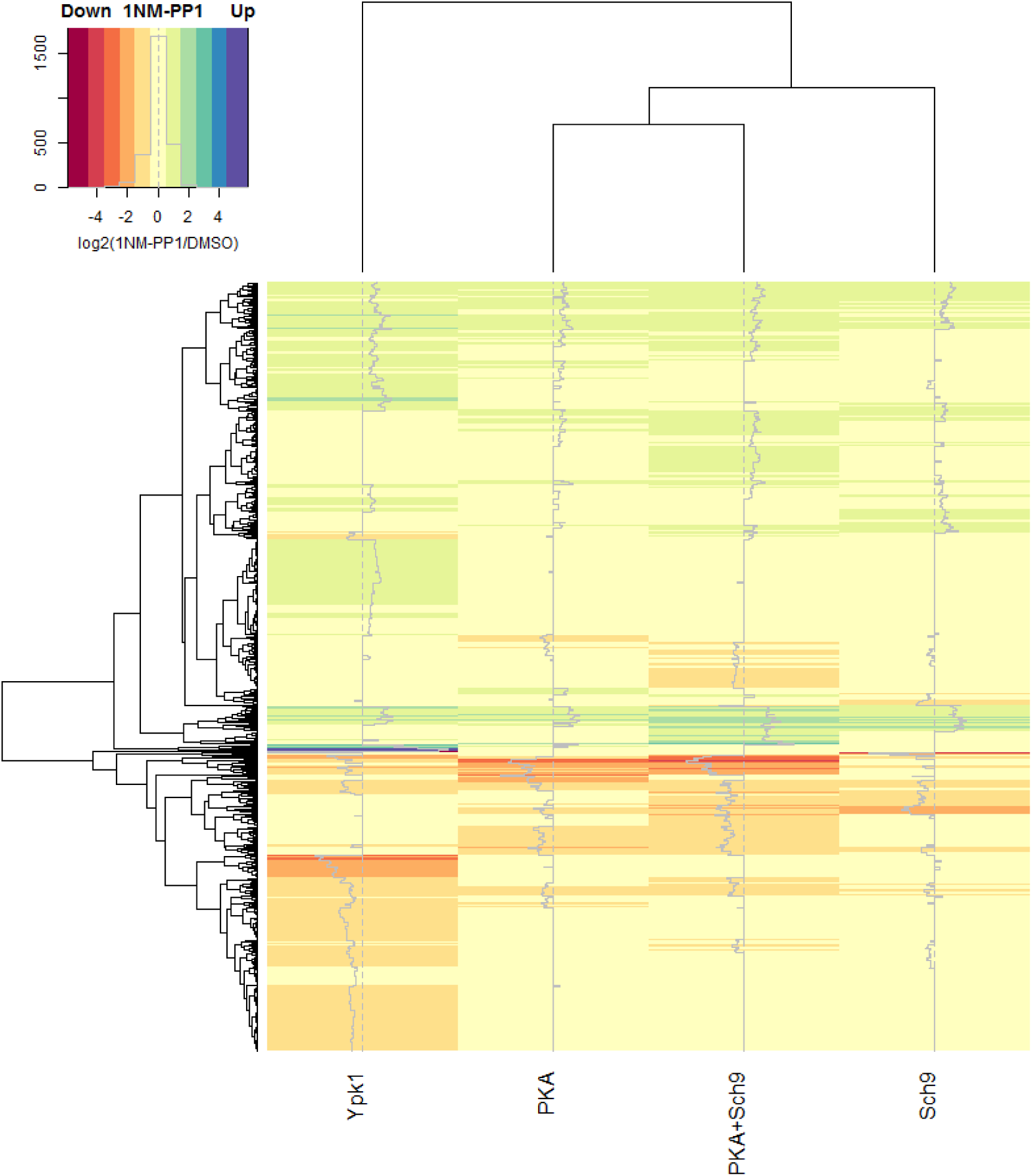
Heatmap of phosphorylation changes upon AGC-kinase inhibition. Mean values for three replicates of log_2_-fold changes in phospho-peptide intensities for 1NM-PP1 vs. DMSO treatment are indicated by color. Data of phospho-peptides significantly (p_Adj_ < 0.05) hyper- or hypo-phosphorylated upon inhibition of at least one kinase in the PKA+Sch9, PKA, Sch9 or Ypk1 dataset are shown.

**Fig. S4.**
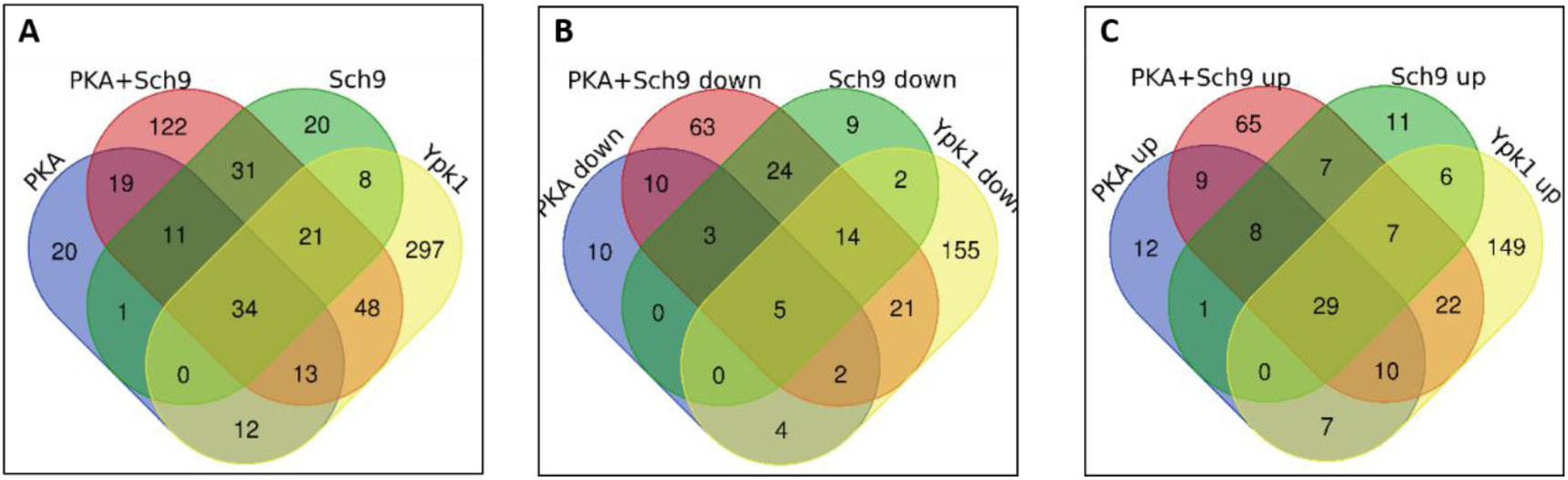
Overlap of phospho-peptides affected by inhibition of different AGC-kinases. Venn diagrams depict phospho-peptides significantly (p_Adj_ < 0.05) affected by inhibition of the kinases indicated. A, Union of hyper- and hypo-phosphorylated phospho-peptides. B, Hypo-phosphorylated phospho-peptides. C, Hyper-phosphorylated phospho-peptides.

**Fig. S5:**
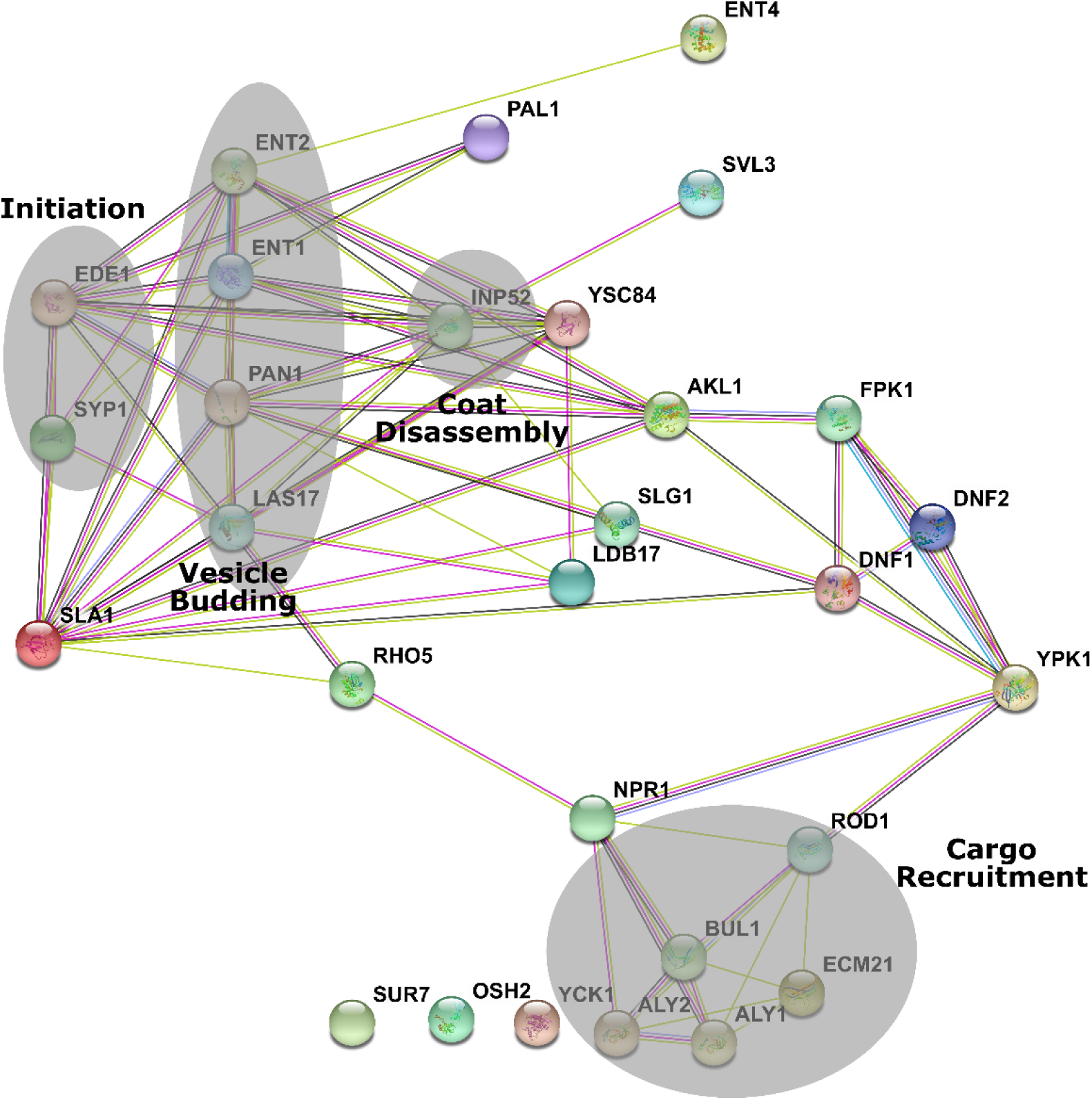
Network of endocytotic proteins affected by Ypk1 inhibition. Proteins with phospho-sites significantly (p < 0.05) affected by Ypk1 inhibition that are associated with the enriched GO-term “Endocytosis” are displayed as a STRING network map. Proteins associated with initiation of endocytosis, vesicle budding, coat disassembly and cargo recruitment are highlighted.

**Fig. S6:**
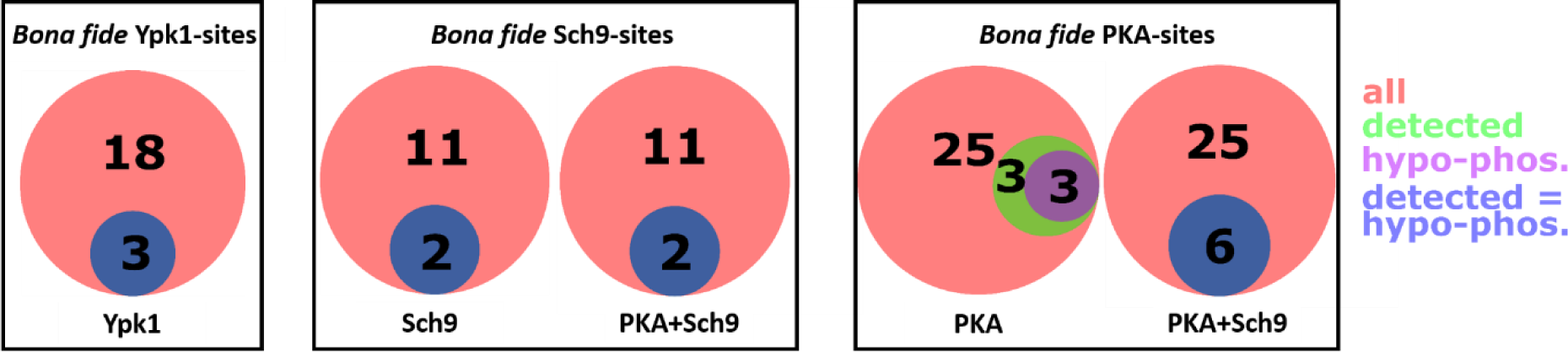
Detected *bona fide* kinase target sites are largely found hypo-phosphorylated upon kinase inhibition. Venn diagrams depicting the number of *bona fide* target sites of the kinase indicated (red), subset of these sites detected by phospho-proteomics (green) and subset of these exhibiting significant (p_Adj_ < 0.05) hypo-phosphorylation (purple). Blue circles indicate a complete overlap of the two subsets.

**Fig. S7:**
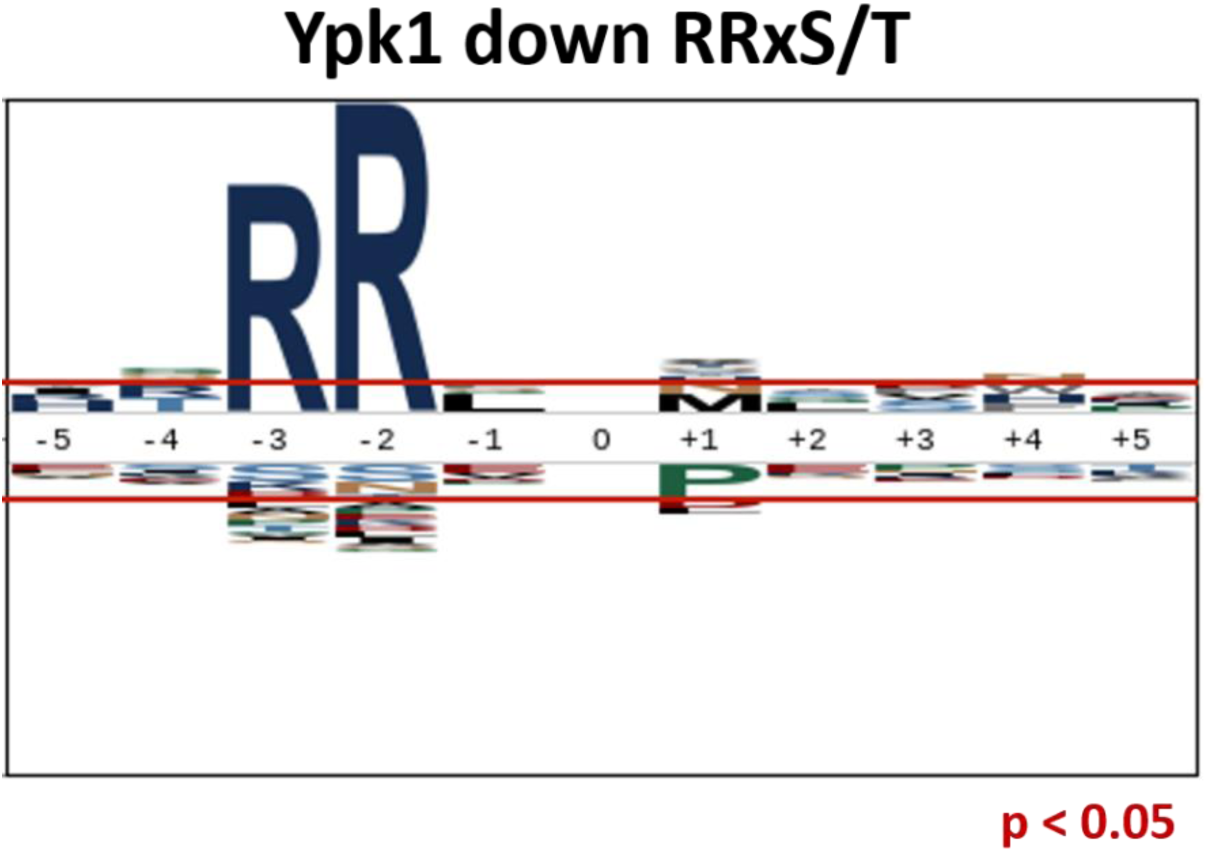
No additional amino acids are enriched around RRxS/T phospho-sites hypo-phosphorylated upon Ypk1 inhibition. A sequence logo was generated from position -5 to +5 around phosphorylated residues using all RRxS/T phospho-sites hypo-phosphorylated upon Ypk1 inhibition as foreground and all quantified phospho-sites as background. Red horizontal lines indicate over- and underrepresentation at p = 0.05.

**Fig. S8:**
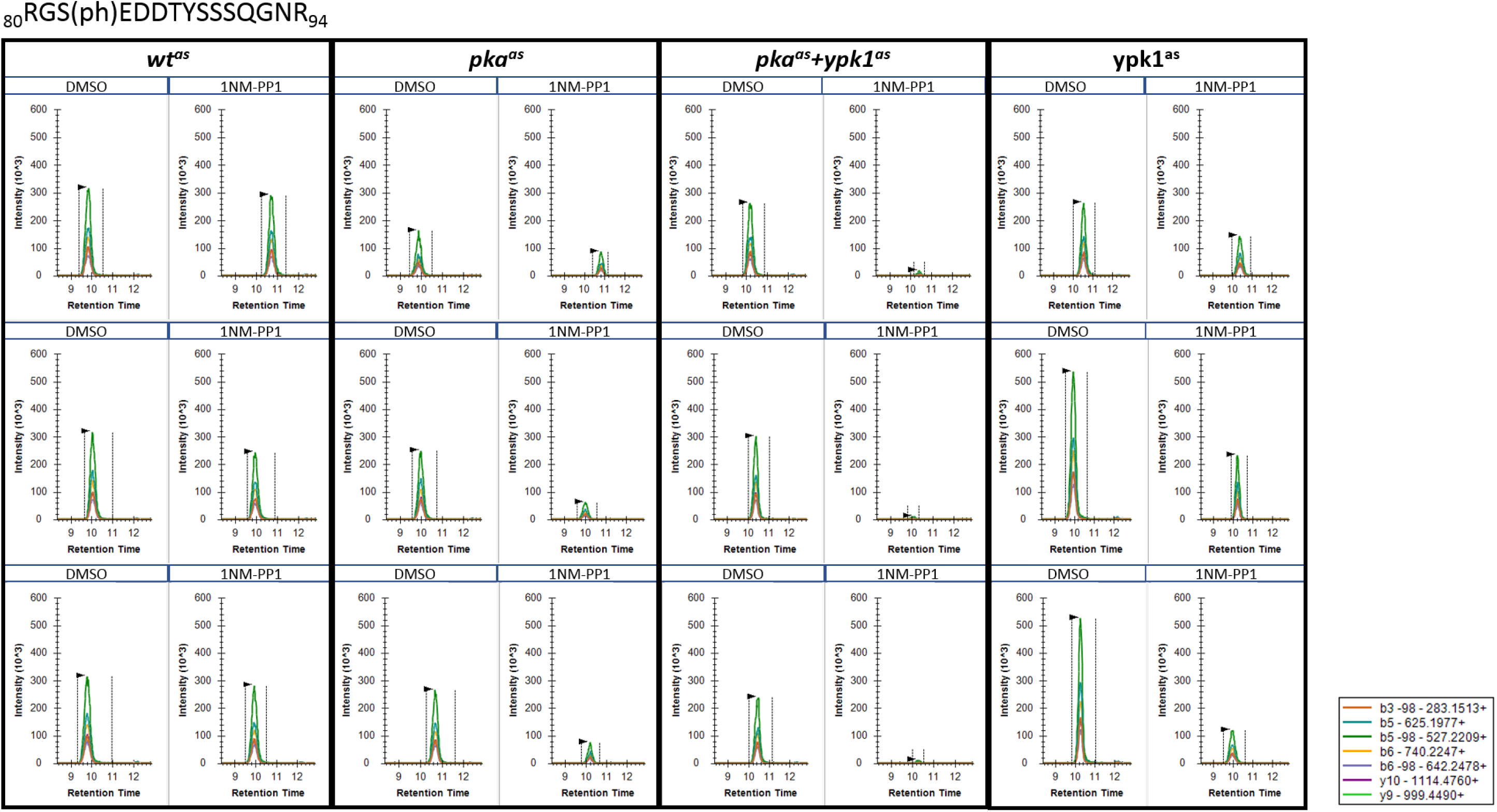
Phosphorylation of Nth1 S83 is reduced upon PKA, Ypk1 and PKA+Ypk1 inhibition. Extracted ion chromatograms of transitions of seven site-determining ions (i.e. ions not shared with other potential phospho-isoforms) of _80_RGS(ph)EDDTYSSSQGNR_94_ (precursor 580.2270; z=3) were extracted from a PRM-experiment on phospho-peptides from strains treated with DMSO or 500 nM 1NM-PP1 for 15 min. Three replicates were analysed per condition.

**Figure S9:**
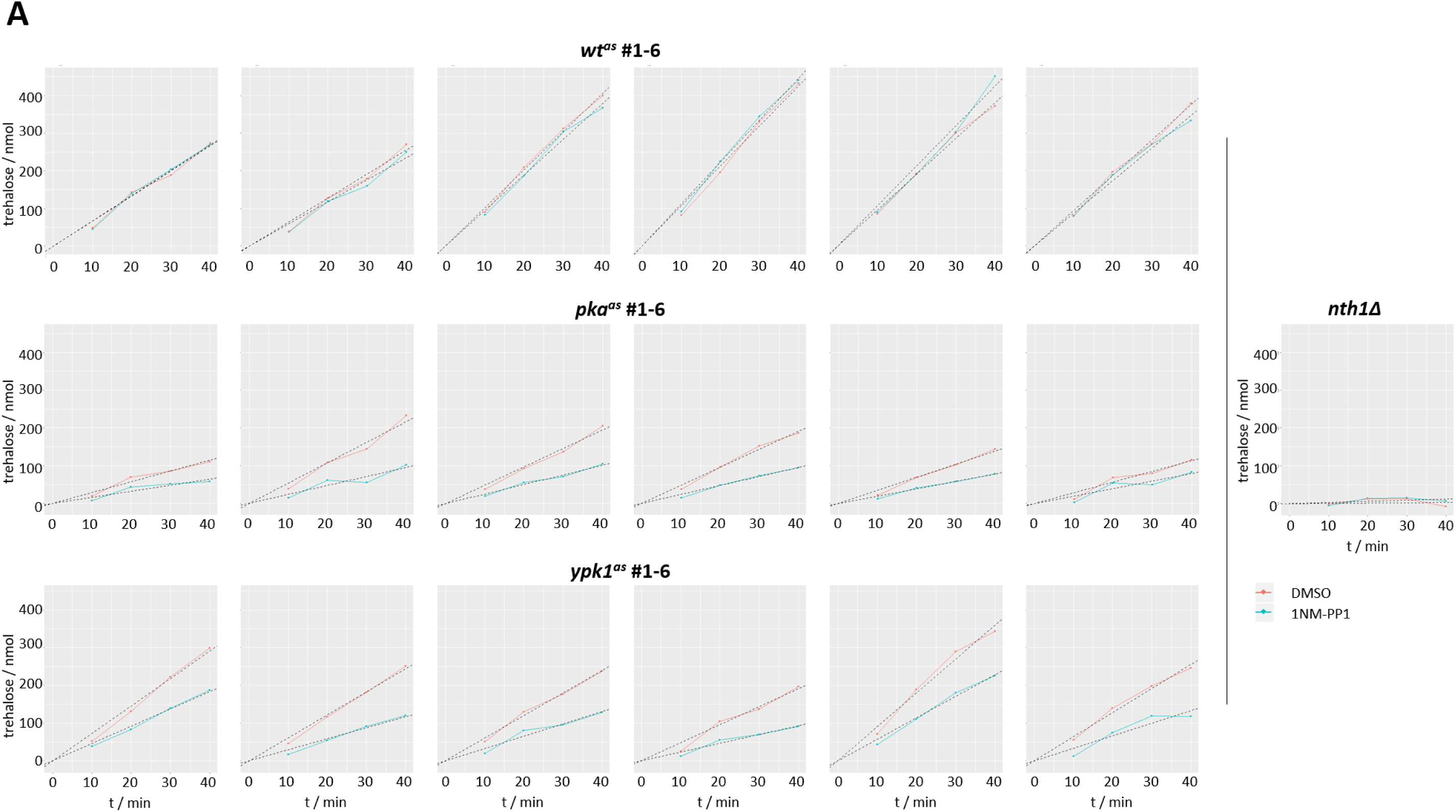
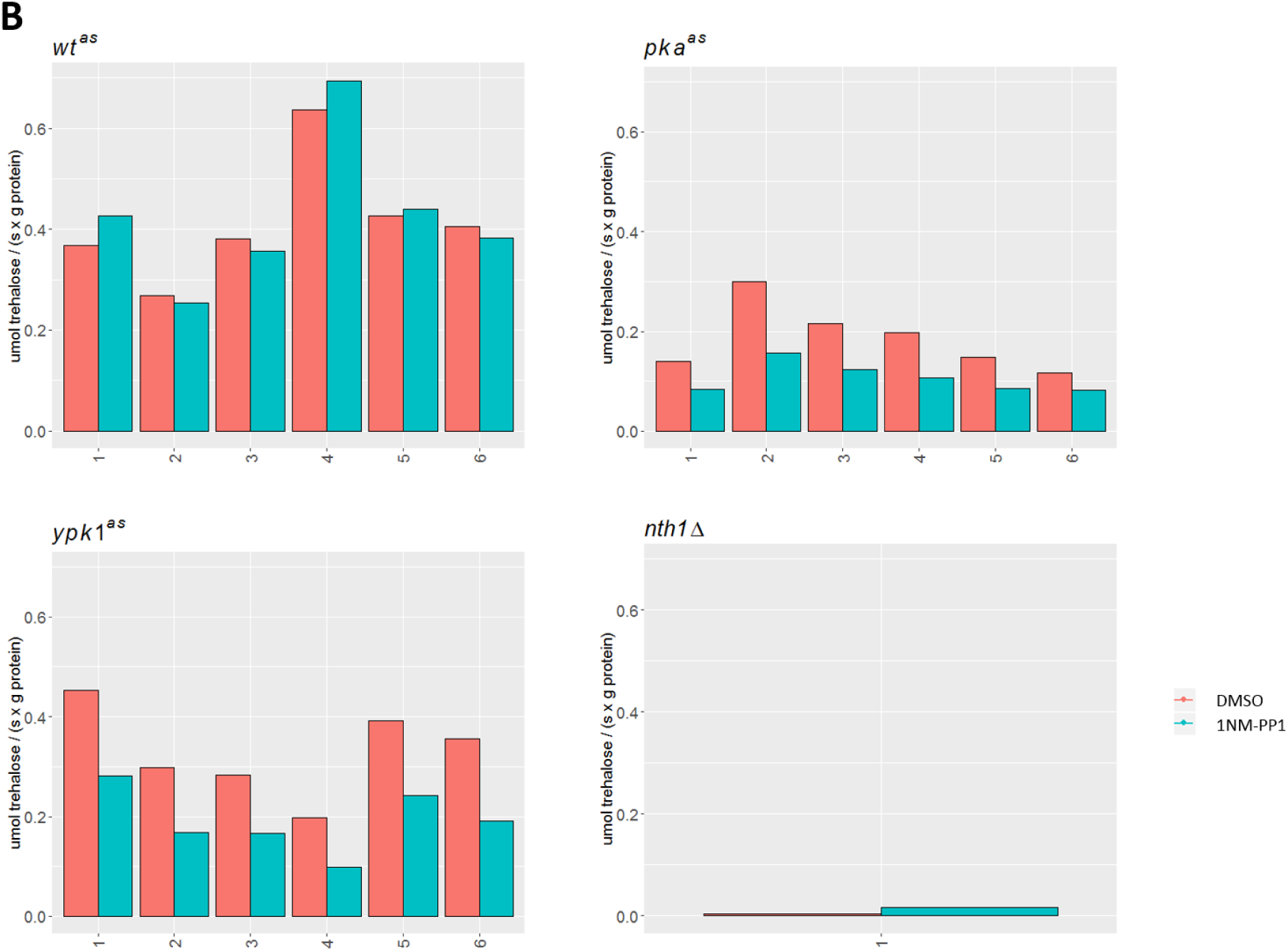
Inhibition of PKA and Ypk1 leads to reduced Nth1 activity. Results of trehalase assay depicted in Figure F. A, Line graphs depicting amount of trehalose consumed over time after addition of trehalose to permeabilized cell pellets of DMSO (red) or 1NM-PP1 (turquoise) treated *wt^as^*, *pka^as^*or *ypk1^as^* (top to bottom) cultures. Graphs for each of the six replicates for these strains are shown. Dashed lines depict linear fit, the slope of which was used to determine specific trehalase activity. Far right: Corresponding plot for *nth1Δ*-strain. B, Specific trehalase activity depicting μmol of trehalose consumed per second and per gram of protein is shown as bar plots for the cultures in A.

**Figure S10:**
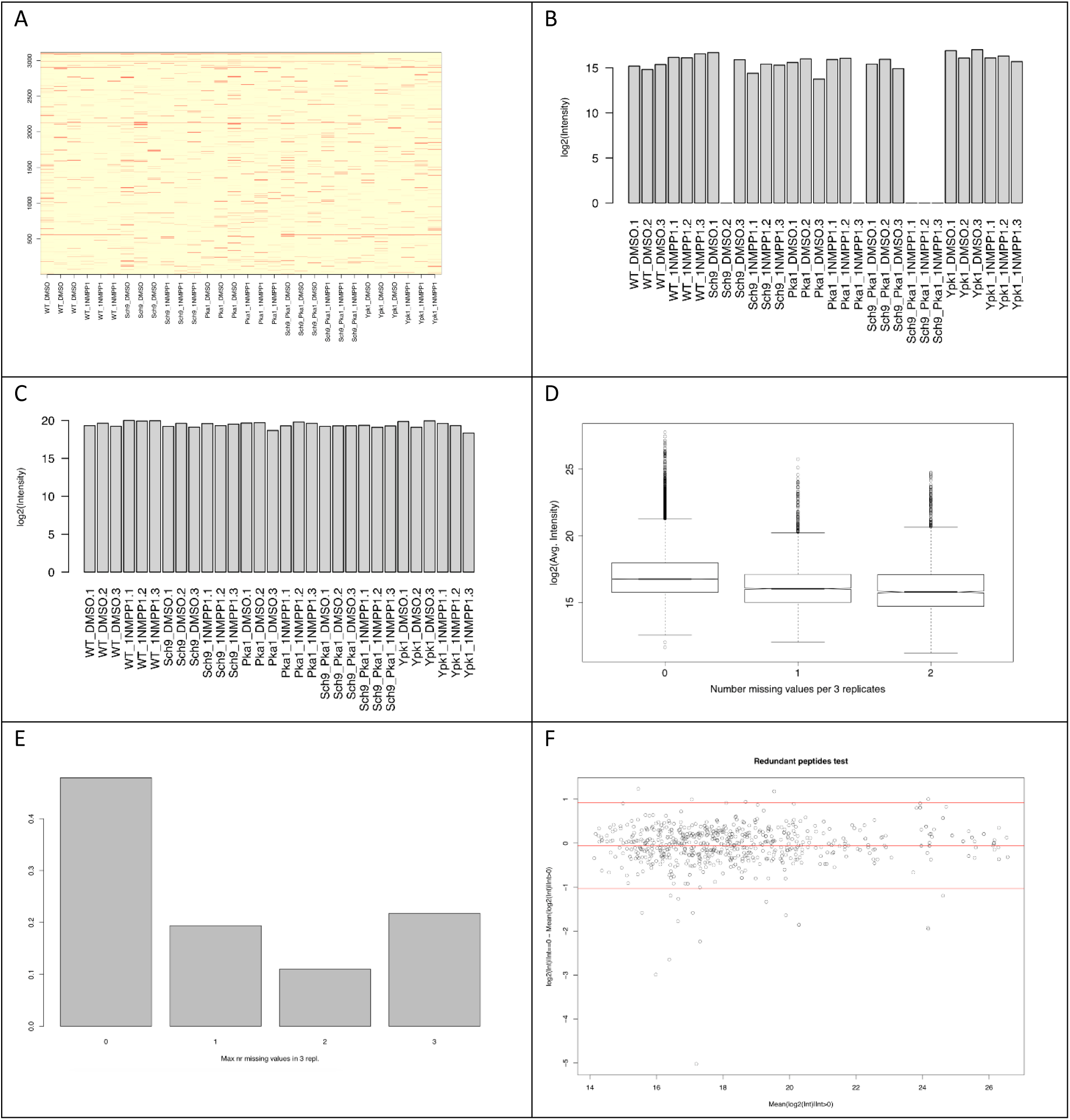
Missing values and their imputation in phospho-proteomics data. A) The occurrence of missing values across the 3000+ phospho-peptides. B) log2-intensity profile of peptide HSS(ph)PDPYGINDKFFDLEK and C) sibling peptide HSS(ph)PDPYGINDK. D) Average log2-intensity of the non-missing values in replicates with 0, 1 and 2 missing values. E) The x-axis shows the maximum number of missing values per 3 replicates in all samples. This barplot displays the frequency of phospho-peptides with a maximum number of 0, 1, 2, and 3 missing values. F) The x-axis is the *α_P’,S,i_* value of all sibling peptides *P*’ without missing values. The y-axis is the difference *d* = *I_P’,S,i_* - *α_P’,S,i_* between the real and estimated value at replicate *i* and sample *S* of *P*’. The horizontal red lines indicate the mean *μ* and *μ±2σ* of the distribution of differences *d*.

## References

1. Loewith R, Hall MN. Target of rapamycin (TOR) in nutrient signaling and growth control. Genetics. 2011 Dec;189(4):1177–201.

2. Urban J, Soulard A, Huber A, Lippman S, Mukhopadhyay D, Deloche O, et al. Sch9 Is a Major Target of TORC1 in Saccharomyces cerevisiae. Mol Cell. 2007 Jun;26(5):663–74.

3. Broach JR. Nutritional control of growth and development in yeast. Genetics. 2012 Sep;192(1):73–105.

4. Schmelzle T, Beck T, Martin DE, Hall MN. Activation of the RAS/cyclic AMP pathway suppresses a TOR deficiency in yeast. Mol Cell Biol. 2004 Jan;24(1):338–51.

5. Zaman S, Lippman SI, Schneper L, Slonim N, Broach JR. Glucose regulates transcription in yeast through a network of signaling pathways. Mol Syst Biol. 2009;5:245.

6. Toda T, Cameron S, Sass P, Wigler M. SCH9, a gene of Saccharomyces cerevisiae that encodes a protein distinct from, but functionally and structurally related to, cAMP-dependent protein kinase catalytic subunits. Genes Dev. 1988 May;2(5):517–27.

7. Broach JR. RAS genes in Saccharomyces cerevisiae: signal transduction in search of a pathway. Trends Genet TIG. 1991 Jan;7(1):28–33.

8. Toda T, Cameron S, Sass P, Zoller M, Wigler M. Three different genes in S. cerevisiae encode the catalytic subunits of the cAMP-dependent protein kinase. Cell. 1987 Jul 17;50(2):277–87.

9. Peeters T, Louwet W, Gelade R, Nauwelaers D, Thevelein JM, Versele M. Kelch-repeat proteins interacting with the G protein Gpa2 bypass adenylate cyclase for direct regulation of protein kinase A in yeast. Proc Natl Acad Sci. 2006 Aug 29;103(35):13034–9.

10. Smith A, Ward MP, Garrett S. Yeast PKA represses Msn2p/Msn4p-dependent gene expression to regulate growth, stress response and glycogen accumulation. EMBO J. 1998 Jul 1;17(13):3556–64.

11. Shenhar G, Kassir Y. A positive regulator of mitosis, Sok2, functions as a negative regulator of meiosis in Saccharomyces cerevisiae. Mol Cell Biol. 2001 Mar;21(5):1603–12.

12. Longo VD. The Ras and Sch9 pathways regulate stress resistance and longevity. Exp Gerontol. 2003 Jul;38(7):807–11.

13. Van de Velde S, Thevelein JM. Cyclic AMP-protein kinase A and Snf1 signaling mechanisms underlie the superior potency of sucrose for induction of filamentation in Saccharomyces cerevisiae. Eukaryot Cell. 2008 Feb;7(2):286–93.

14. Kamada Y, Fujioka Y, Suzuki NN, Inagaki F, Wullschleger S, Loewith R, et al. Tor2 Directly Phosphorylates the AGC Kinase Ypk2 To Regulate Actin Polarization. Mol Cell Biol. 2005 Aug 15;25(16):7239–48.

15. Niles BJ, Mogri H, Hill A, Vlahakis A, Powers T. Plasma membrane recruitment and activation of the AGC kinase Ypk1 is mediated by target of rapamycin complex 2 (TORC2) and its effector proteins Slm1 and Slm2. Proc Natl Acad Sci. 2012 Jan 31;109(5):1536–41.

16. Chen P, Lee KS, Levin DE. A pair of putative protein kinase genes (YPK1 and YPK2) is required for cell growth in Saccharomyces cerevisiae. Mol Gen Genet MGG. 1993 Jan;236(2–3):443–7.

17. Berchtold D, Piccolis M, Chiaruttini N, Riezman I, Riezman H, Roux A, et al. Plasma membrane stress induces relocalization of Slm proteins and activation of TORC2 to promote sphingolipid synthesis. Nat Cell Biol. 2012 Apr 15;14(5):542–7.

18. Niles BJ, Powers T. TOR complex 2–Ypk1 signaling regulates actin polarization via reactive oxygen species. Mol Biol Cell. 2014;25(24):3962–3972.

19. Hatano T, Morigasaki S, Tatebe H, Ikeda K, Shiozaki K. Fission yeast Ryh1 GTPase activates TOR Complex 2 in response to glucose. Cell Cycle Georget Tex. 2015;14(6):848–56.

20. Eltschinger S, Loewith R. TOR Complexes and the Maintenance of Cellular Homeostasis. Trends Cell Biol. 2016 Feb;26(2):148–59.

21. Casamayor A, Torrance PD, Kobayashi T, Thorner J, Alessi DR. Functional counterparts of mammalian protein kinases PDK1 and SGK in budding yeast. Curr Biol CB. 1999 Feb 25;9(4):186–97.

22. Bodenmiller B, Aebersold R. Quantitative analysis of protein phosphorylation on a system-wide scale by mass spectrometry-based proteomics. Methods Enzymol. 2010;470:317–34.

23. Bishop AC, Buzko O, Shokat KM. Magic bullets for protein kinases. Trends Cell Biol. 2001 Apr;11(4):167–72.

24. van der Plaat JB. Cyclic 3’,5’-adenosine monophosphate stimulates trehalose degradation in baker’s yeast. Biochem Biophys Res Commun. 1974 Feb 4;56(3):580–7.

25. Schepers W, Van Zeebroeck G, Pinkse M, Verhaert P, Thevelein JM. In vivo phosphorylation of Ser21 and Ser83 during nutrient-induced activation of the yeast protein kinase A (PKA) target trehalase. J Biol Chem. 2012 Dec 28;287(53):44130–42.

26. Panek A. Synthesis of trehalose by baker’s yeast (Saccharomyces cerevisiae). Arch Biochem Biophys. 1962 Sep;98:349–55.

27. Singer MA, Lindquist S. Multiple effects of trehalose on protein folding in vitro and in vivo. Mol Cell. 1998 Apr;1(5):639–48.

28. van Heerden JH, Wortel MT, Bruggeman FJ, Heijnen JJ, Bollen YJM, Planqué R, et al. Lost in transition: start-up of glycolysis yields subpopulations of nongrowing cells. Science. 2014 Feb 28;343(6174):1245114.

29. Wera S, De Schrijver E, Geyskens I, Nwaka S, Thevelein JM. Opposite roles of trehalase activity in heat-shock recovery and heat-shock survival in Saccharomyces cerevisiae. Biochem J. 1999 Nov 1;343 Pt 3:621–6.

30. Jorgensen P, Rupes I, Sharom JR, Schneper L, Broach JR, Tyers M. A dynamic transcriptional network communicates growth potential to ribosome synthesis and critical cell size. Genes Dev. 2004 Oct 15;18(20):2491–505.

31. Yorimitsu T, Zaman S, Broach JR, Klionsky DJ. Protein kinase A and Sch9 cooperatively regulate induction of autophagy in Saccharomyces cerevisiae. Mol Biol Cell. 2007 Oct;18(10):4180–9.

32. Huber A, Bodenmiller B, Uotila A, Stahl M, Wanka S, Gerrits B, et al. Characterization of the rapamycin-sensitive phosphoproteome reveals that Sch9 is a central coordinator of protein synthesis. Genes Dev. 2009 Aug 15;23(16):1929–43.

33. deHart AKA, Schnell JD, Allen DA, Hicke L. The conserved Pkh-Ypk kinase cascade is required for endocytosis in yeast. J Cell Biol. 2002 Jan 21;156(2):241–8.

34. Kaksonen M, Roux A. Mechanisms of clathrin-mediated endocytosis. Nat Rev Mol Cell Biol. 2018 May;19(5):313–26.

35. Schmidt A, Beck T, Koller A, Kunz J, Hall MN. The TOR nutrient signalling pathway phosphorylates NPR1 and inhibits turnover of the tryptophan permease. EMBO J. 1998 Dec 1;17(23):6924–31.

36. Kennelly PJ, Krebs EG. Consensus sequences as substrate specificity determinants for protein kinases and protein phosphatases. J Biol Chem. 1991 Aug 25;266(24):15555–8.

37. Mok J, Kim PM, Lam HYK, Piccirillo S, Zhou X, Jeschke GR, et al. Deciphering protein kinase specificity through large-scale analysis of yeast phosphorylation site motifs. Sci Signal. 2010 Feb 16;3(109):ra12.

38. Muir A, Ramachandran S, Roelants FM, Timmons G, Thorner J. TORC2-dependent protein kinase Ypk1 phosphorylates ceramide synthase to stimulate synthesis of complex sphingolipids. eLife. 2014 Oct 3;3.

39. Roelants FM, Baltz AG, Trott AE, Fereres S, Thorner J. A protein kinase network regulates the function of aminophospholipid flippases. Proc Natl Acad Sci. 2010 Jan 5;107(1):34–9.

40. Roelants FM, Breslow DK, Muir A, Weissman JS, Thorner J. Protein kinase Ypk1 phosphorylates regulatory proteins Orm1 and Orm2 to control sphingolipid homeostasis in Saccharomyces cerevisiae. Proc Natl Acad Sci. 2011 Nov 29;108(48):19222–7.

41. Muir A, Roelants FM, Timmons G, Leskoske KL, Thorner J. Down-regulation of TORC2-Ypk1 signaling promotes MAPK-independent survival under hyperosmotic stress. eLife. 2015 Aug 14;4.

42. Huber A, French SL, Tekotte H, Yerlikaya S, Stahl M, Perepelkina MP, et al. Sch9 regulates ribosome biogenesis via Stb3, Dot6 and Tod6 and the histone deacetylase complex RPD3L. EMBO J. 2011 Jul 5;30(15):3052–64.

43. Moir RD, Lee J, Haeusler RA, Desai N, Engelke DR, Willis IM. Protein kinase A regulates RNA polymerase III transcription through the nuclear localization of Maf1. Proc Natl Acad Sci U S A. 2006 Oct 10;103(41):15044–9.

44. Reinders A, Bürckert N, Boller T, Wiemken A, De Virgilio C. Saccharomyces cerevisiae cAMP-dependent protein kinase controls entry into stationary phase through the Rim15p protein kinase. Genes Dev. 1998 Sep 15;12(18):2943–55.

45. Roelants FM, Leskoske KL, Martinez Marshall MN, Locke MN, Thorner J. The TORC2-Dependent Signaling Network in the Yeast Saccharomyces cerevisiae. Biomolecules. 2017 05;7(3).

46. Kuret J, Johnson KE, Nicolette C, Zoller MJ. Mutagenesis of the regulatory subunit of yeast cAMP-dependent protein kinase. Isolation of site-directed mutants with altered binding affinity for catalytic subunit. J Biol Chem. 1988 Jul 5;263(19):9149–54.

47. Ma P, Wera S, Van Dijck P, Thevelein JM. The PDE1-encoded low-affinity phosphodiesterase in the yeast Saccharomyces cerevisiae has a specific function in controlling agonist-induced cAMP signaling. Mol Biol Cell. 1999 Jan;10(1):91–104.

48. Nash P, Tang X, Orlicky S, Chen Q, Gertler FB, Mendenhall MD, et al. Multisite phosphorylation of a CDK inhibitor sets a threshold for the onset of DNA replication. Nature. 2001 Nov 29;414(6863):514–21.

49. Alblova M, Smidova A, Docekal V, Vesely J, Herman P, Obsilova V, et al. Molecular basis of the 14-3-3 protein-dependent activation of yeast neutral trehalase Nth1. Proc Natl Acad Sci U S A. 2017 14;114(46):E9811–20.

50. Veisova D, Macakova E, Rezabkova L, Sulc M, Vacha P, Sychrova H, et al. Role of individual phosphorylation sites for the 14-3-3-protein-dependent activation of yeast neutral trehalase Nth1. Biochem J. 2012 May 1;443(3):663–70.

51. Lee YJ, Jeschke GR, Roelants FM, Thorner J, Turk BE. Reciprocal Phosphorylation of Yeast Glycerol-3-Phosphate Dehydrogenases in Adaptation to Distinct Types of Stress. Mol Cell Biol. 2012 Nov 15;32(22):4705–17.

52. François J, Parrou JL. Reserve carbohydrates metabolism in the yeast Saccharomyces cerevisiae. FEMS Microbiol Rev. 2001 Jan;25(1):125–45.

53. De Virgilio C, Bürckert N, Boller T, Wiemken A. A method to study the rapid phosphorylation-related modulation of neutral trehalase activity by temperature shifts in yeast. FEBS Lett. 1991 Oct 21;291(2):355–8.

54. Budhwar R, Lu A, Hirsch JP. Nutrient control of yeast PKA activity involves opposing effects on phosphorylation of the Bcy1 regulatory subunit. Mol Biol Cell. 2010 Nov 1;21(21):3749–58.

55. Lorenz MC, Pan X, Harashima T, Cardenas ME, Xue Y, Hirsch JP, et al. The G protein-coupled receptor gpr1 is a nutrient sensor that regulates pseudohyphal differentiation in Saccharomyces cerevisiae. Genetics. 2000 Feb;154(2):609–22.

56. Schmitz H-P, Jendretzki A, Sterk C, Heinisch JJ. The Small Yeast GTPase Rho5 and Its Dimeric GEF Dck1/Lmo1 Respond to Glucose Starvation. Int J Mol Sci. 2018 Jul 26;19(8).

57. Kraakman L, Lemaire K, Ma P, Teunissen AW, Donaton MC, Van Dijck P, et al. A Saccharomyces cerevisiae G-protein coupled receptor, Gpr1, is specifically required for glucose activation of the cAMP pathway during the transition to growth on glucose. Mol Microbiol. 1999 Jun;32(5):1002–12.

58. Xue Y, Batlle M, Hirsch JP. GPR1 encodes a putative G protein-coupled receptor that associates with the Gpa2p Galpha subunit and functions in a Ras-independent pathway. EMBO J. 1998 Apr 1;17(7):1996–2007.

59. Ptacek J, Devgan G, Michaud G, Zhu H, Zhu X, Fasolo J, et al. Global analysis of protein phosphorylation in yeast. Nature. 2005 Dec 1;438(7068):679–84.

60. Galello F, Portela P, Moreno S, Rossi S. Characterization of substrates that have a differential effect on Saccharomyces cerevisiae protein kinase A holoenzyme activation. J Biol Chem. 2010 Sep 24;285(39):29770–9.

61. De Virgilio C, Loewith R. Cell growth control: little eukaryotes make big contributions. Oncogene. 2006 Oct 16;25(48):6392–415.

62. Alvaro CG, Aindow A, Thorner J. Differential Phosphorylation Provides a Switch to Control How α-Arrestin Rod1 Down-regulates Mating Pheromone Response in Saccharomyces cerevisiae. Genetics. 2016;203(1):299–317.

63. Amberg DC, Burke D, Strathern JN, Burke D. Methods in yeast genetics: a Cold Spring Harbor Laboratory course manual. 2005 ed. Cold Spring Harbor, N.Y: Cold Spring Harbor Laboratory Press; 2005. 230 p.

64. Cox J, Mann M. MaxQuant enables high peptide identification rates, individualized p.p.b.-range mass accuracies and proteome-wide protein quantification. Nat Biotechnol. 2008 Dec;26(12):1367–72.

65. Cox J, Neuhauser N, Michalski A, Scheltema RA, Olsen JV, Mann M. Andromeda: a peptide search engine integrated into the MaxQuant environment. J Proteome Res. 2011 Apr 1;10(4):1794–805.

66. Smyth GK. Linear models and empirical bayes methods for assessing differential expression in microarray experiments. Stat Appl Genet Mol Biol. 2004;3:Article3.

67. O’Shea JP, Chou MF, Quader SA, Ryan JK, Church GM, Schwartz D. pLogo: a probabilistic approach to visualizing sequence motifs. Nat Methods. 2013 Dec;10(12):1211–2.

68. Cherry JM, Hong EL, Amundsen C, Balakrishnan R, Binkley G, Chan ET, et al. Saccharomyces Genome Database: the genomics resource of budding yeast. Nucleic Acids Res. 2012 Jan;40(Database issue):D700–705.

69. Snel B, Lehmann G, Bork P, Huynen MA. STRING: a web-server to retrieve and display the repeatedly occurring neighbourhood of a gene. Nucleic Acids Res. 2000 Sep 15;28(18):3442–4.

70. MacLean B, Tomazela DM, Shulman N, Chambers M, Finney GL, Frewen B, et al. Skyline: an open source document editor for creating and analyzing targeted proteomics experiments. Bioinforma Oxf Engl. 2010 Apr 1;26(7):966–8.

